# Global Mapping of Combinatorial Chromatin Regulatory Events Using Hi-Plex CUT&Tag

**DOI:** 10.1101/2025.10.06.680180

**Authors:** Yuan Liao, Weiqiang Zhou, Min-Zhi Jiang, Kuai Yu, Zixiao Jolene Wang, Thomas Drummond, Meng Ding, Yingming Zhao, Ignacio Pino, Sean D. Taverna, Hongkai Ji, Heng Zhu

## Abstract

Epigenetic modulators, transcription factors, and chromatin-associated proteins collaboratively regulate essential genomic functions, such as transcription, repression, and DNA damage repair. However, the intricate cross-talk or the interaction between these chromatin regulatory factors (CRFs) remains poorly understood, largely due to limitations in experimental methods, which traditionally interrogate the genomic locations of only one CRF. Although the latest methods can study multiple CRFs simultaneously, they cannot examine the interaction between these factors. Inferring biological interactions often requires integrating data from multiple experiments, which can be labor-intensive and imprecise since the measurement is not from the same DNA molecule. To address these challenges, we developed Hi-Plex CUT&Tag, an advanced technology that enables the robust detection of co-localization events across over 600 pairs of CRFs using 36 barcoded monoclonal antibodies (mAbs), while minimizing background signals and cross-contamination. Hi-Plex CUT&Tag facilitates comprehensive pairwise analysis of epigenetic modifiers, including histone post-translational modifications (PTMs), writers, and transcription factors (TFs). Each Tn5 tagmentation fragment provides detailed insights, capturing the identities of two co-localized events, the underlying genomic sequences, and the molecular distance between them. For the first time, numerous novel bivalent events, epigenetic-context-dependent transcriptional regulation, and specific chromatin mark combinations with significant impacts on gene regulation can be detected in a single experiment. Furthermore, single-cell Hi-Plex CUT&Tag enables the analysis of synergistic interactions between CRF pairs at the single-cell level, providing unprecedented resolution for studying chromatin dynamics.

## INTRODUCTION

In eukaryotic cells, DNA is wrapped around histone proteins, forming nucleosomes, the fundamental subunits of chromatin. Approximately 30 million nucleosomes are distributed throughout the chromatin of most human cells, however, the functional properties of these nucleosomes vary widely across the genome. Although we are only beginning to understand the molecular basis for this functional heterogeneity, post-translational modifications (PTMs) on histones are known to chemically alter the local chromatin environment. These modifications can serve as binding platforms and influence nucleosome structure for chromatin-associated complexes that regulate gene transcription and otherwise modulate the functional state of chromatin. The hypothesis that histone PTMs act individually or in combination at the level of individual nucleosomes is often referred to as the “epigenetic code” or “histone code”. This code has the potential to greatly expand the informational capacity of the genetic code, providing deeper insights into the regulation and function of chromatin [1–4]. Beyond histone PTMs, the enzymes that write, erase, and read these modifications, along with the multi-protein complexes they form, and transcription factors (TFs), also work together in a coordinated and dynamic manner to regulate the accessibility and function of chromatin-associated proteins along the genome. Without a comprehensive understanding of how these events co-localize in space and time, it remains challenging to make robust predictions about gene expression and cellular phenotypes.

A fundamental hurdle in deciphering the molecular cross-talk that regulates chromatin-based biological processes lies in the technical challenges of probing how histone PTMs, chromatin regulators, and TFs, collectively referred to as chromatin regulatory factors (CRFs), co-localize on small stretches of nucleosomes. Integrated analyses from two decades of protein and genome-wide sequencing studies suggest that dozens of distinct histone PTM combinations may co-occur on a single nucleosome. However, most *in vivo* epigenetic profiling methods, including ChIP-seq [5–7], CUT&RUN [8], and CUT&Tag [9], are limited to measuring one type of event at a time, such as a specific histone PTM, epigenetic modifier, or TF. Consequently, co-localization is often inferred by aligning data from separate experiments, which do not measure the same cell and DNA molecule, and may be performed under different conditions or by different research groups. These limitations leave the combinatorial interactions among histone marks and chromatin-associated complexes on the same DNA molecule or chromatin allele largely unexplored, impeding a deeper understanding of their coordinated roles in gene regulation.

Recently, several methods, such as MulTI-Tag [10] and CoTarget [11], have been developed to achieve multiplexed functionality by performing multiple consecutive rounds of CUT&Tag. MulTI-Tag [10] and MAbID [12] further improve specificity by conjugating barcoded adapters with antibodies. Another multiplexed method, ChIP-DIP [13], combines barcodes and antibodies on protein G beads to profile hundreds of targets simultaneously. However, none of these methods can directly detect co-localized CRFs on the same DNA molecule within the same cell, limiting their ability to capture the spatial and functional interactions critical for understanding chromatin dynamics.

Methods that enable multiplexed mapping of multiple CRFs within the same cell have been developed. For example, Tn5 transposase-based approaches, such as Multi-CUT&Tag [14, 15], NTT-seq [16], and Nano-CT [17, 18], utilize barcoded adapters preloaded onto Tn5 transposase to profile multiple targets simultaneously. While these methods may capture co-localization of paired CRFs within the same cell, the weak affinity between Protein A and IgG can cause Tn5 to detach and engage in open chromatin tagmentation, leading to high background noise. This issue becomes more pronounced when multiple antibody-Protein A-Tn5 complexes are mixed, as pre-loaded DNA barcodes can also detach from Tn5 and exchange between complexes, generating significant cross-contamination signals [10]. These limitations restrict their multiplexing capacity to a small number of monoclonal antibodies (mAbs), thereby constraining their utility for high-plex high-resolution chromatin studies.

To address these limitations, we present a novel technology, Hi-Plex CUT&Tag, which enables simultaneous, pairwise genome-wide profiling of multiple CRFs by significantly reducing background noise and cross-contamination. By multiplexing 36 mAbs within the same cells, this approach allows us to globally analyze co-localization events across over 600 CRF pairs, with detailed information about their genomic locations, allelic distribution, and relative spacing. For the first time, these datasets have enabled us to identify and map genome-wide, numerous poorly studied DNA-associated events co-occurring across the same small stretch of chromatin, investigate epigenetic context-dependent transcriptional regulation, and discern specific CRF combinations with significant regulatory roles.

Additionally, we have applied Hi-Plex CUT&Tag to single cells, which facilitates the examination of synergistic and antagonistic interactions among different chromatin mark pairs by analyzing their co-occurrence patterns within individual cells. This innovation provides unprecedented resolution for understanding the dynamic interplay of chromatin marks in regulating gene expression and chromatin organization.

We believe that the Hi-Plex CUT&Tag approach represents a significant advancement in chromatin profiling. Its improved specificity, sensitivity, and ability to detect multiple targets with a streamlined workflow and precise control over the tagmentation process make it a powerful tool for studying chromatin biology, interplays among histone PTMs, histone modification enzymes and DNA-binding proteins, and epigenetic cross-talk in diverse biological contexts.

## RESULTS

### Hi-Plex CUT&Tag enables concurrent genome-wide profiling of multiple CRFs

Hi-Plex CUT&Tag advances multiplex chromatin profiling by enabling simultaneous genome-wide mapping of multiple CRFs, including histone PTMs, histone-modifying enzymes, and DNA-binding proteins (DBPs) such as TFs, as well as their pairwise co-localization events on the same DNA molecule or chromatin allele (**Fig. 1A**). To enable highly multiplexed tagmentation via reducing background signals and cross-contamination, we developed a novel strategy for barcoding mAbs and the tagmentation process. Using tetrameric streptavidin as a connector, we conjugated biotinylated, barcoded DNA adapter sequences to each biotinylated mAb individually before pooling them. The nearly irreversible biotin-streptavidin interaction (*K_D_* < 10⁻¹⁴ M) [19] ensures that the DNA adapter sequences remain stably attached to the mAbs, preventing dissociation and minimizing non-specific ATAC-like background signals or cross-contamination caused by adapter DNA swapping. Additionally, we introduced unloaded Tn5 and MgCl₂ only after unbound mAbs were removed from the cells, further reducing background noise. This modification also better preserves the activity of Tn5 by eliminating the need for overnight incubation.

**Figure 1.**
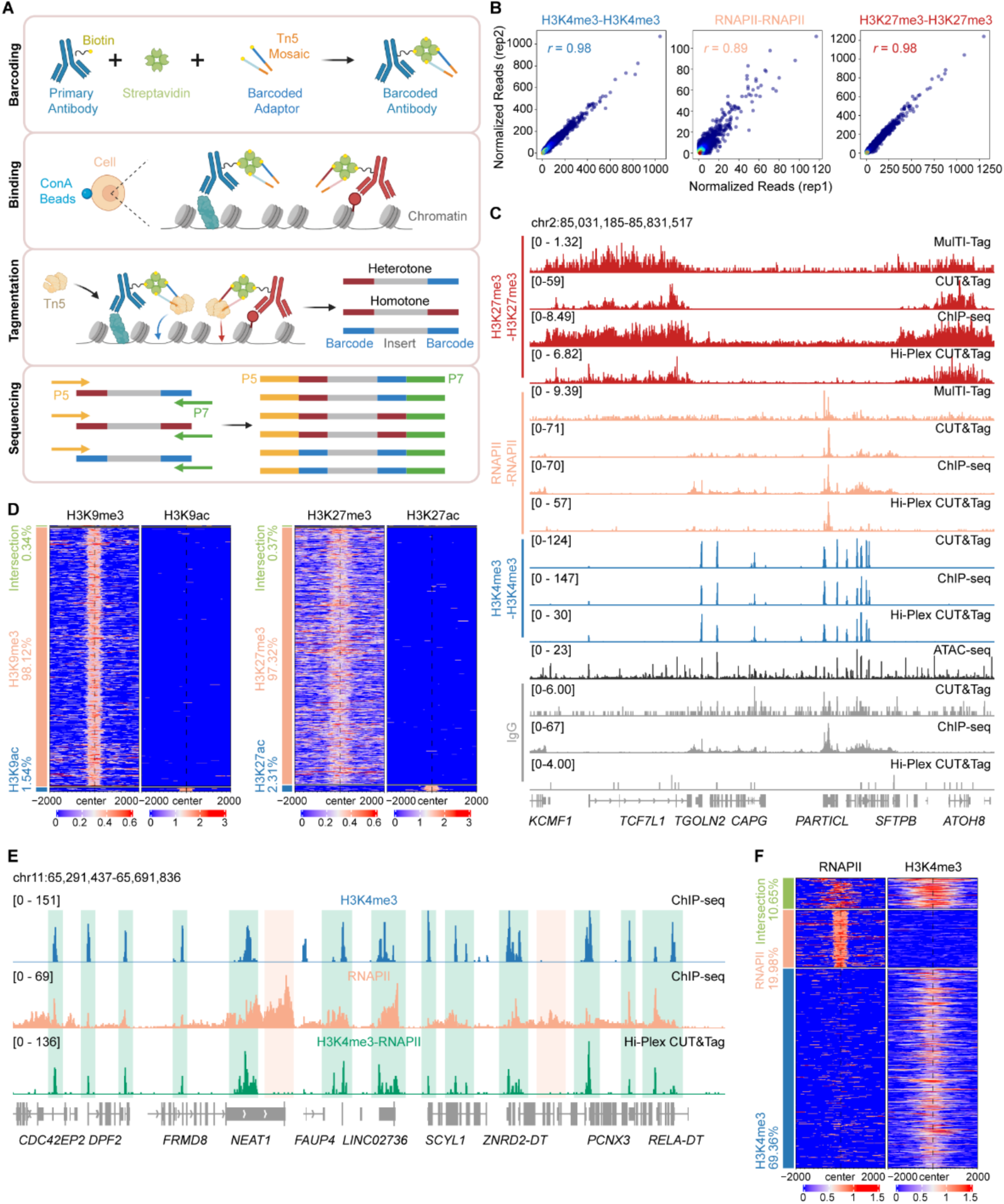
Concurrent and effective characterization of multiple chromatin proteins using Hi-Plex CUT&Tag. **A**. Hi-Plex CUT&Tag workflow. 1. Primary antibodies are individually biotinylated and conjugated with adaptor DNA sequences, containing Tn5 binding mosaic (orange), specific barcode sequences and P5 or P7 sequence (blue), via streptavidin. 2. Individual barcoded antibodies are pooled and incubated with permeabilized cells immobilized on beads. 3. After unbound antibodies are removed, Tn5 and MgCl_2_ are added to tagment nearby genomic DNA. The two ends of a chromosomal DNA fragment can be tagged with two different barcodes (i.e., heterotone) or the same barcodes (i.e., homotone). 4. Fragment libraries are enriched by PCR and sequenced by Illumina NGS. **B**. Scatter plot analysis confirms high reproducibility between the two replicates, as illustrated by H3K4me3, RNAPII and H3K27me3 homotone data. *r*: Pearson correlation. **C**. Comparison of signal tracks of H3K27me3, RNAPII, H3K4me3 homotone and IgG singletone reads between Hi-Plex CUT&Tag, MulTI-Tag, CUT&Tag and ChIP-seq in RPM (reads per million), as well as ATAC-seq obtained in K562. Hi-Plex CUT&Tag profiles showed a similar profile to those obtained with the other methods, suggesting comparable coverage of this new technology. Note that although the track of euchromatin mark H3K4me3 showed substantial overlap with those obtained by ATAC-seq, additional ATAC signals are found in regions covered by H3K27me3, indicating that Hi-Plex signals are less likely to be contaminated by random open chromatin signals. Hi-Plex CUT&Tag has the lowest IgG background signal. **D**. Comparison between mutually exclusive histone marks: H3K9me3 homotone versus H3K9ac homotone and H3K27me3 homotone versus H3K27ac homotone. Almost no overlapping signals are found on the chromatins between these exclusive marks. **E**. Examples of signal comparison between the H3K4me3–RNAPII heterotone track and individual H3K4me3 and RNAPII ChIP-seq tracks. Green shaded areas highlight peaks where Hi-Plex CUT&Tag heterotone signals overlap with both individual ChIP-seq peaks. In contrast, the orange shaded area indicates regions where H3K4me3–RNAPII heterotone peaks are missing, corresponding to the absence of one individual ChIP-seq signal. **F**. Heatmaps showing substantial, global overlapping signals between RNAPII and H3K4me3 homotone peaks.

To demonstrate the multiplexing capacity and accuracy of Hi-Plex CUT&Tag, we barcoded 36 mAbs, targeting 12 common histone marks, 14 histone-modifying enzymes, 8 human TFs, CTCF, and phoshorylated Ser2 of PolII (RNAPII) (**Supplementary Fig. S1C**). A rabbit IgG was included as a negative control. The 37 barcoded antibodies were pooled and incubated with permeabilized K562 cells (10⁵ cells) for 1 hour at room temperature. After stringent washes, unloaded Tn5 and MgCl₂ were added for tagmentation at 37 °C for 1 hour. The genomic DNA was then extracted, PCR-amplified, and sequenced (**Fig. 1A**). This experiment was performed in duplicate, yielding ∼200 million reads.

After de-multiplexing, 92.6% of the reads were successfully assigned to their respective antibodies. When two different barcode sequences representing two distinct CRFs are found on the ends of a tagmented sequence, we classify it as a heterotone event. In contrast, when the same barcode sequences are found on the ends of a tagmented sequence, we classify it as a homotone event (**Fig. 1A**).

To evaluate the assay reproducibility, we compared the data of replicated assays and found high correlations between the replicates, indicating robustness of the method (**Fig. 1B**). Comparative analyses demonstrated a high degree of similarity between Hi-Plex CUT&Tag homotone data and corresponding MulTI-Tag, CUT&Tag or ChIP-seq data. For instance, the H3K27me3, RNAPII, and H3K4me3 profiles generated by Hi-Plex CUT&Tag closely matched those previously obtained with MulTI-Tag, CUT&Tag, and ENCODE ChIP-seq (**Fig. 1C**). To further validate the dataset, we compared it with existing ATAC-seq data. We observed that nearly all H3K4me3 homotone peaks were contained by the ATAC-seq peaks. However, many additional ATAC-seq peaks were present, indicating that Hi-Plex CUT&Tag peaks are antibody-specific and not influenced by non-specific open chromatin signals (**Fig. 1C**).

To illustrate the specificity of Hi-Plex CUT&Tag, we selected pairs of mutually exclusive histone marks and compared their homotone data. For example, consistent with the repressive role of H3K27me3, Hi-Plex CUT&Tag demonstrated mutually exclusive signals between H3K27me3 homotone and RNAPII homotone at most genomic regions (**Fig. 1C**). In addition, minimum overlapping homotone reads were found between other mutually exclusive marks, such as H3K9me3 vs. H3K9ac and H3K27me3 vs. H3K27ac, further confirming the low cross-contamination between antibodies (**Fig. 1D**).

To assess background noise, we pooled all homotone and heterotone reads containing the rabbit IgG barcode, creating a pooled dataset. Because the pooled reads share only a single barcode, they are referred to as IgG singleton data, analogous to a CUT&Tag or traditional ChIP-seq dataset (**Supplementary Fig. S1A**). We found that the IgG singleton dataset only accounted for 0.07% of the total reads, indicating minimal background. Comparing our IgG singletone data to the counterparts obtained with ChIP-seq and CUT&Tag in the same cell line [9, 20], we observed substantially lower IgG signals with Hi-Plex CUT&Tag in regions with comparable chromatin mark signals (**Fig. 1C**).

Together, these results demonstrate that Hi-Plex CUT&Tag generates antibody-specific, low-background signals and effectively profiles multiple CRFs with high sensitivity, specificity, and reproducibility.

### Hi-Plex CUT&Tag enables mapping of co-localized CRF pairs on the same DNA molecule

Each tagmentation generated by Hi-Plex CUT&Tag requires two proximal tagmentation events on the same chromatin in the same cell within a certain distance (approximately 1 kb) because no adapter sequences are used. Each tagmentation corresponds to a pair of distinct CRFs (i.e., heterotone) or the same CRF (i.e., homotone), and the fragment length approximates the distance between the two CRFs. Distinct DNA barcodes are used to represent different CRFs recognized by the corresponding mAbs, allowing sequencing reads, especially the heterotone reads, to carry pairwise co-localization information of CRFs. Hi-Plex CUT&Tag thus provides data along three axes: CRF pair, distance between CRFs (reflected by DNA fragment size), and genomic location of the event (**Fig. 2A**).

**Figure 2.**
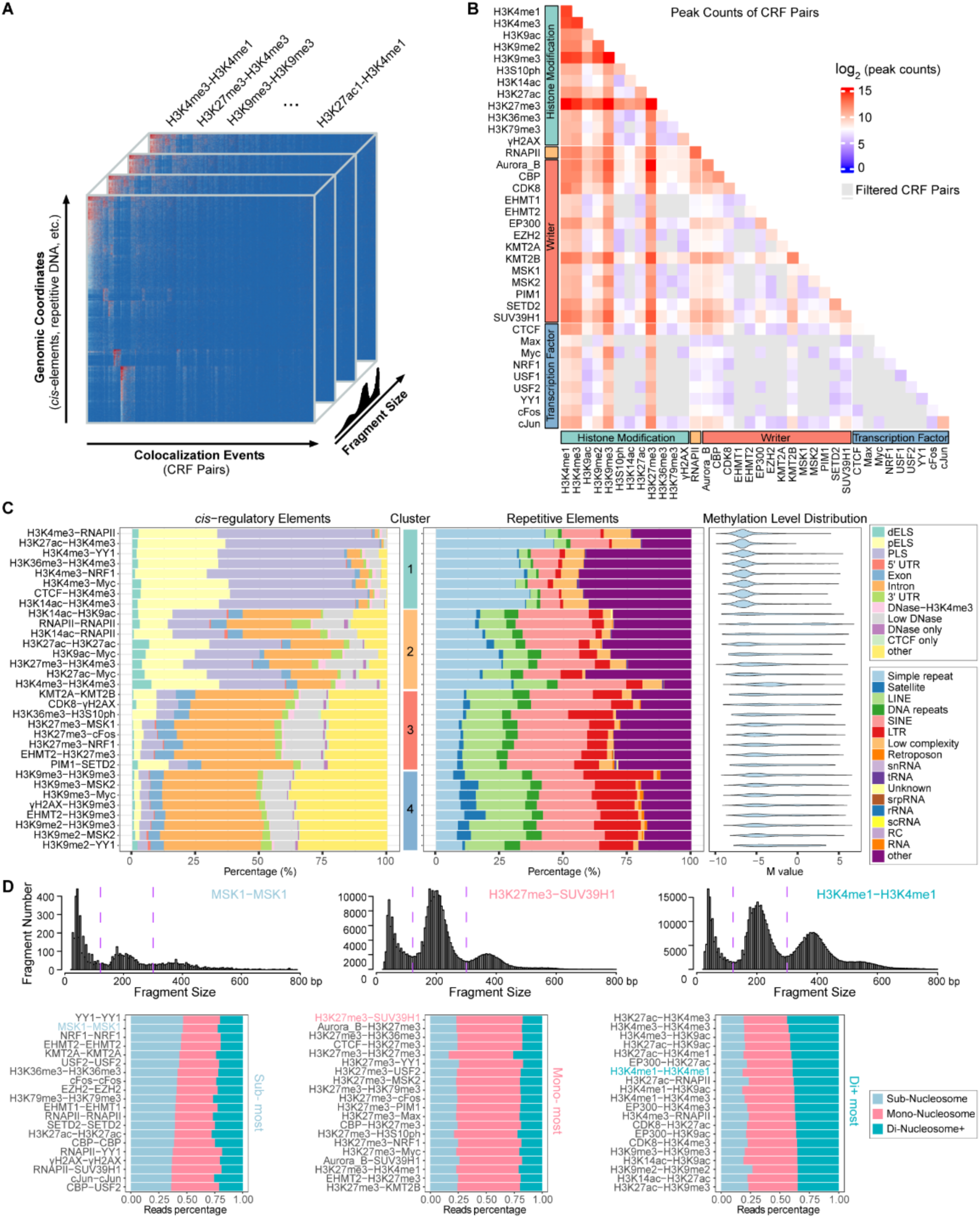
Dissecting the complexity of Hi-Plex CUT&Tag data. **A**. Illustration of the information packed in the sequencing reads obtained with Hi-Plex CUT&Tag technology. This technology can simultaneously map where the co-localization of hundreds of CRF pairs occur on the whole genome and provide the distance between each co-localized CRF pair. Various regulatory or repetitive elements can also be readily annotated to those sequencing reads. **B**. Summary of numbers of peaks of 630 heterotone and 36 homotone pairs in a heatmap. CRF pairs that did not pass the cutoff value are labeled in grey and removed from further analyses. **C**. Peak region annotation of each CRF pair represented by stacked bar plots using the ENCODE candidate *cis*-regulatory elements (cCREs), repetitive elements, and DNA methylation, respectively. Four major clusters are identified using hierarchical clustering based on the *cis*-regulatory elements profiles of the CRF pairs. Eight representative CRF pairs from each cluster were shown. **D.** Top: Histograms of fragment length distributions for three example CRF pairs. DNA fragments are classified into sub-nucleosome, mono-nucleosome, and di+-nucleosome categories, with cutoffs indicated by dashed vertical lines. Bottom: Stacked bar charts of fragment length distributions for the top 20 ranked CRF pairs, ordered by the proportion of reads in each category. One representative example from each category is shown in the top histograms. CRF pairs connecting two transcription factors tend to yield shorter fragments (sub-nucleosome), whereas CRF pairs associated with euchromatin marks more often generate longer fragments (di+-nucleosome).

By our definition, a heterotone read is generated by two distinct CRFs (e.g., H3K4me3 and RNAPII), unambiguously reflecting the co-localization of two CRFs on the same chromosomal fragment within the same cell (**Fig. 1A**). A homotone event, on the other hand, may or may not genuinely represent the co-localization of two CRFs of the same type, especially for sequences shorter than 300 bp, as a very small fraction of the barcoded mAbs carried more than one pair of the Tn5 adapter sequences, capable of generating a homotone read by a single CRF (**Supplementary Fig. S1B**). Among all mapped reads, approximately 68% were classified as heterotone events, suggesting that Hi-Plex CUT&Tag could effectively detect co-localization of CRF pairs.

To validate the ability of Hi-Plex CUT&Tag to detect CRF co-localization, we compared Hi-Plex CUT&Tag heterotone data to ChIP-seq data of corresponding histone PTMs obtained in the same K562 cell line. For example, heterotone tracks of transcription-associated H3K4me3 and RNAPII were compared to ENCODE ChIP-seq tracks for these marks. H3K4me3-RNAPII heterotone tracks overlapped regions where H3K4me3 and RNAPII ChIP-seq signals co-occurred (green shaded areas; **Fig. 1E**). Conversely, regions lacking H3K4me3 ChIP-seq signals, such as the *NEAT1* gene body, also lacked H3K4me3-RNAPII heterotone signals (orange shaded areas; **Fig. 1E**). A global comparison between the homotone data of the two CRFs further confirmed overlapping signals between these two marks, supporting the accuracy of Hi-Plex CUT&Tag in detecting co-localization events (**Fig. 1F**).

### A systematic characterization of co-localized CRF pairs

Each fragment generated by Hi-Plex CUT&Tag contains rich information, including the identities of the two co-localized CRFs, the underlying genomic sequences where the CRF pairs were positioned, and the molecular distance between the CRFs or cut sites (**Fig. 2A**). Using 36 barcoded mAbs, Hi-Plex CUT&Tag could presumably examine 36 homotone and 630 (=36x35/2) heterotone CRF pairs. Analysis in K562 cells revealed variability in the number of reads and peaks across CRF pairs, likely reflecting differences in CRF abundance, CRF availability, distance between CRFs, and/or CRF stability (**Fig. 2B**). For example, H3K4me1, H3K4me3, H3K9me3, and H3K27me3 mAbs produced the most reads and peaks, followed by proteins, such as RNAPII, Aurora B, CBP, CDK8, EP300, KMT2B, SETD2, and SUV39H1. Fewer reads were generated for most TFs tested, likely due to the lower frequency of TF-DNA binding events compared to the more abundant histone PTMs along chromatin. Our lack of using chromatin cross-linking, to preserve CRF accessibility, may also reduce TF stability on the chromatin. After filtering out CRF pairs with low quality (**Methods**), 501 CRF pairs were retained for further analysis.

To characterize the epigenetic roles of CRF pairs, their peaks were annotated using the ENCODE Registry of Candidate *Cis*-Regulatory Elements (cCREs) [21] (**Supplementary Table S1**), repetitive elements [22] (**Supplementary Table S1**), and DNA methylation data [20] (**Methods**). The 501 CRF pairs are grouped into four major clusters based on their cCREs annotation (**Fig. 2C** - Eight representative ; see **Supplementary Fig. S2D** for the comprehensive clustering).

Different clusters exhibit distinct genomic and epigenetic characteristics. Cluster 1 is dominated by euchromatin marks, including CRF pairs such as H3K4me3-RNAPII and H3K27ac-H3K4me3 (**Fig. 2C**). Peaks in this cluster are highly enriched for proximal enhancer-like (pELS) and promoter-like (PLS) elements. The cluster also shows a high proportion of simple repeats and low transposon content, such as LINE, SINE, and LTR.

Cluster 2 consists of CRF pairs involving RNAPII and histone acetylation marks, such as H3K14ac-RNAPII and RNAPII-RNAPII, as well as enhancer marks, such as H3K27ac (**Fig. 2C**). This cluster has an increased proportion of peaks located in gene body (i.e., exons, introns, and UTRs) and distal enhancer-like elements (dELS). The enrichment of gene body features in RNAPII and histone acetylation marks suggests a role in transcriptional elongation, because RNA polymerase II actively transcribes genes and acetylation is associated with an open chromatin structure, facilitating efficient transcription. The cluster also exhibits increased proportions of SINE and LINE repetitive elements. Indeed, previous studies have shown that K562 cells express full-length L1 mRNAs and L1-encoded proteins [23, 24].

Cluster 3 contains many CRF pairs involving H3K27me3, indicating repressive functions (**Fig. 2C**). Peaks are enriched in gene body and low-DNase regions. Repetitive elements are mostly dominated by SINE, LINE, and LTR.

Cluster 4 features CRF pairs involving H3K9me2 and H3K9me3, associated with heterochromatin (**Fig. 2C**). Peaks are enriched for gene body and low-DNase regions. The repetitive elements are mostly SINE, LINE and LTR, and showed an increased proportion of satellite DNA, a major component of centromeric and pericentromeric DNA.

The DNA methylation levels at the peak regions of CRF pairs in each cluster reflect their regulatory roles. Clusters 1 and 2, containing mostly open chromatin marks, showed relatively lower DNA methylation levels, whereas clusters 3 and 4, associated with repressive marks, exhibited broader DNA methylation distributions, consistent with the role of CpG methylation in gene silencing [25] (**Fig. 2C**).

Interestingly, we observed that tagmented sequences displayed a distinct laddering pattern, suggesting structural periodicity (**Supplementary Fig. S2A**). Fragment length distribution analysis revealed three major peaks at approximately 60, 200, and 380 bp, separated by valleys in between them, which correspond respectively to sub-, mono-, di- and weakly discernible tri+ nucleosome fragments (**Supplementary Fig. S2B**). We ranked CRF pairs based on the percentages of sub-, mono-, di+-nucleosome DNA fragments, respectively, with examples of top pairs in each category shown in **Fig. 2D**. Notably, sub-nucleosome fragments (<80 bp) are most enriched for CRF pairs involving TFs, such as YY1, NRF1, cFos, and USF2, reflecting their short sequence-specific binding footprints in the linker regions between nucleosomes. On the other hand, the top-ranked CRF pairs enriched for di+-nucleosome fragments (>300 bp) involve pairs between euchromatin histone marks (e.g., H3K27ac-H3K4me3) and/or their writers (e.g., EP300-H3K27ac and EP300-H3K9ac). The longer fragments may represent previously proposed mechanisms of chromatin modifiers “reading and writing” histone modifications across several nucleosomes. Interestingly, most of the top-ranked CRF pairs enriched for mono-nucleosome fragments involve H3K27me3, which may reflect its high density in heterochromatin.

### Hi-Plex CUT&Tag enables comprehensive analysis of combinatorial landscape of colocalized CRF pairs

The concurrent profiling of 501 CRF pairs within the same sample enables comprehensive examination of combinatorial co-occurrence pattern of CRF pairs within the same genomic region, which is not feasible using traditional ChIP-seq, CUT&RUN, or CUT&Tag approaches. To leverage this unique capability and better understand the relationship between CRFs and transcriptional regulation, we divided the genome into 800 bp non-overlapping regions and characterized each region using the 501 CRF pairs. After filtering out regions that lack substantial signals and selecting the highly variable regions across the CRF pairs, we performed unsupervised clustering analysis on the remaining 105,519 genomic regions (covering 84.4 million base pairs), identifying 16 CRF pair clusters (indexed 1–16) and 15 genomic region clusters (indexed A–O) (**Methods**). A subset of representative regions is shown in **Fig. 3A**, and the full set of regions is shown in **Supplementary Fig. S3A**.

**Figure 3.**
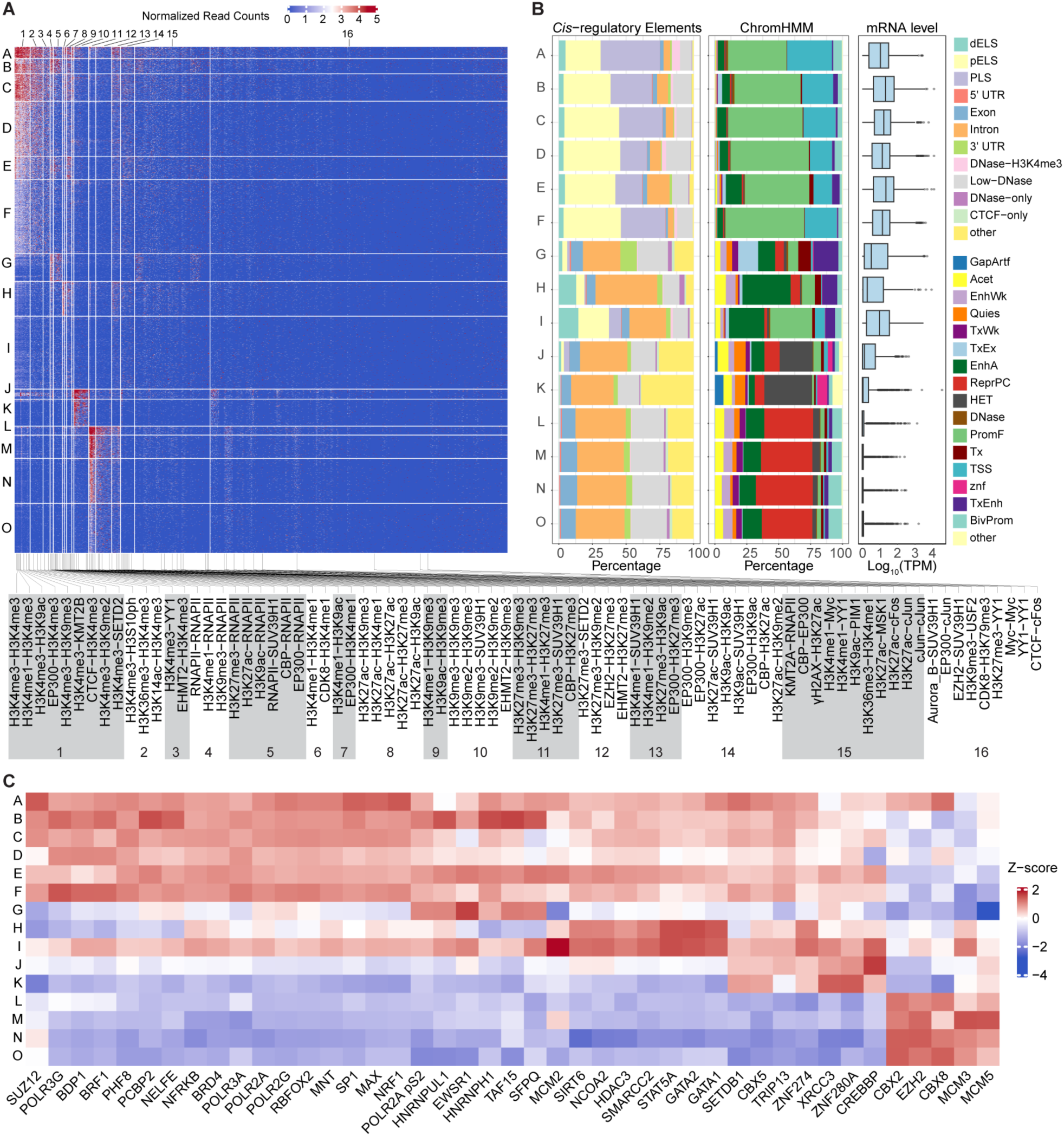
Hi-Plex CUT&Tag reveals the combinatorial landscape of co-localized CRF pairs. **A.** Biclustering analysis of K562 Hi-Plex CUT&Tag data across 501 CRF pairs identified 16 CRF pair clusters (1–16) and 15 genomic region clusters (A–O), each displaying distinct combinatorial co-localization patterns. Heatmap colors indicate normalized signals for each CRF pair in the corresponding genomic regions (blue = low, red = high). Representative CRF pairs from each cluster are shown below the heatmap. **B**. Genomic regions in each cluster were annotated using ENCODE candidate *cis*-regulatory elements and ChromHMM states. Stacked bar plots indicate annotation categories (colored), and RNA-seq expression of overlapping genes is shown as side boxplots. **C**. Transcription factor binding site enrichment analysis revealed distinct patterns across region clusters. Heatmap colors represent standardized log-odds ratios (blue = low enrichment, red = high enrichment).

Regarding CRF pair clusters (**Fig. 3A; Supplementary Table S2**), Clusters 1, 2, and 3 consist of CRF pairs involving H3K4me3, a mark for active promoters. Cluster 1 contains H3K4me3 homotone and heterotones paired with other CRFs, including heterochromatin marks (e.g., H3K9me3 and H3K9me2), activation or enhancer marks (e.g., H3K9ac and H3K4me1), CTCF, EP300, SETD2, and KMT2B. Cluster 2 contains H3K4me3 heterotones paired with H3S10ph, H3K36me3, and H3K14ac. Cluster 3 contains H3K4me3 heterotones paired with YY1 and EHMT2. Clusters 4 and 5 consist of CRF pairs involving RNA Polymerase II (RNAPII), which has critical functions in gene transcription. Cluster 4 contains the RNAPII homotone and heterotones that pair RNAPII with H3K4me1 and H3K9me3. Cluster 5 contains heterotones that pair RNAPII with H3K27me3, H3K27ac, H3K9ac, SUV39H1, CBP, and EP300. Clusters 6 and 7 consist of CRF pairs involving H3K4me1, which marks active and poised enhancers and promoters. Cluster 6 contains the H3K4me1 homotone and heterotone between H3K4me1 and CDK8. Cluster 7 contains heterotones that pair H3K4me1 with H3K9ac and EP300. Cluster 8 contains the homotone of the enhancer mark H3K27ac and heterotones that pair H3K27ac with H3K4me1, H3K9ac, H3K9me3, and H3K27me3. Cluster 9 contains heterotone CRF pairs involving H3K9me3, such as H3K4me1-H3K9me3 and H3K9ac-H3K9me3. Cluster 10 consists of homotone pairs involving two heterochromatin marks, H3K9me3 and H3K9me2, as well as heterotones that pair them with other marks such as SUV39H1 and EHMT2. Clusters 11 and 12 consist of CRF pairs involving the heterochromatin mark H3K27me3. Cluster 11 contains the H3K27me3 homotone and heterotones that pair H3K27me3 with H3K9me3, H3K4me1, SUV39H1, and CBP. Cluster 12 contains heterotones that pair H3K27me3 with SETD2, H3K9me2, EZH2, and EHMT2. Cluster 13 contains heterotone CRF pairs involving H3K27me3 (e.g., H3K27me3-H3K9ac, EP300-H3K27me3) or H3K4me1 (e.g. H3K4me1-SUV39H1, H3K4me1-H3K9me2).

Cluster 14 consists of CRF pairs involving acetylation marks H3K27ac, H3K9ac, and acetyltransferases such as EP300 and CBP (e.g., H3K9ac-H3K9ac, EP300-H3K9ac, EP300-H3K27ac, CBP-H3K27ac). It also contains heterotone CRF pairs with potentially opposing functions, such as EP300-H3K9me3, H3K27ac−SUV39H1, H3K9ac−SUV39H1, and H3K27ac−H3K9me2. Clusters 15 and 16 contain CRF pairs with relatively sparser signals. Cluster 15 mostly contains heterotones that pair euchromatin marks such as H3K4me1, H3K27ac, and H3K9ac, RNAPII, and acetyltransferase (CBP and EP300) with less abundant CRFs such as H3K36me3, gH2AX, PIM1, KMT2A, MSK1, and transcription factors (e.g., YY1, cJun, cFos, and Myc). Cluster 16 mostly contains CRF pairs involving repressive chromatin marks such as H3K9me3 and H3K27me3 (e.g., H3K9me3-USF2 and H3K27me3-YY1), Polycomb Repressive Complexes (EZH2 and SUV39H1), transcript factors (e.g., Myc-Myc and YY1-YY1), and CRF pairs involving CTCF.

Different region clusters display distinct combinatorial patterns of CRF co-localization and activities. We annotated each region cluster using ENCODE cCREs [21] and ChromHMM [26] (**Supplementary Table S1**). We also examined the expression levels of genes linked to each cluster using RNA-seq in K562 cells (**Fig. 3B, Methods**).

Genomic regions in Clusters A, B, C, D, E, and F consist of high proportions of proximal enhancer-like and promoter-like elements (pELS and PLS) compared to other clusters based on cCRE annotation, which is also consistent with high proportions of promoter flanking and active transcript start site regions (PromF and TSS) based on ChromHMM annotation (**Fig. 3B**). Genes associated with these clusters showed high expression levels. These region clusters showed high signals in CRF pair Clusters 1 and 2 (**Fig. 3A**), both involving H3K4me3. Within these region clusters, Cluster E showed relatively higher signals in CRF pair Cluster 8 (**Fig. 3A**), which consists of CRF pairs involving H3K27ac, an active enhancer mark, than the other five region clusters. This may explain why genes associated with Cluster E had relatively higher expression levels and more enhancer regions (EnhA) than the other five clusters.

Region Cluster G consists of regions spanning pELS, PLS, Exon, Intron, and 3’ UTR. These genomic regions are highly related to the gene transcription process (**Fig. 3B**). Interestingly, Cluster G showed high signals in CRF pair Clusters 4 and 5 (**Fig. 3A**), which contain CRF pairs involving RNAPII, and some CRF pairs in Cluster 15 (**Fig. 3A**) which also involve RNAPII. The composition of genomic regions in Cluster G reflects the dynamic process of RNAPII during gene transcription, which moves from the promoter toward the 3’ UTR. Genes associated with Cluster G also showed moderate expression levels.

Region Clusters H and I consist of the highest proportions of distal enhancer-like elements (dELS) and enhancers (EnhA), based on the ENCODE cCRE and ChromHMM annotations, respectively (**Fig. 3B**). Cluster I showed higher proportions of proximal enhancer-like (pELS) and promoter elements (PLS or PromF) elements than Cluster H. Cluster H showed relatively higher signals in CRF pair Clusters 6 and 7 than Cluster I (**Fig. 3A**), which consist of CRF pairs involving H3K4me1, a mark for active and poised enhancers. Cluster I showed relatively higher signals in CRF Cluster 1, which contains CRF pairs involving H3K4me3, than Cluster H (**Fig. 3A, Supplementary Fig. S3A**). Interestingly, we observed that genes associated with Cluster I were highly expressed, while genes associated with Cluster H showed moderate gene expression.

Region Clusters J and K consist of high proportions of heterochromatin regions marked by H3K9me3 (HET) as annotated by ChromHMM (**Fig. 3B**). Consistent with the annotation, these two region clusters showed high signals in CRF pair Cluster 10 (**Fig. 3A**) which consists of CRF pairs involving H3K9me3. Genes associated with these two region clusters showed low expression level.

Region Clusters L, M, N, and O consist of the highest proportions of Polycomb repressive regions marked by H3K27me3 (ReprPC) as annotated by ChromHMM (**Fig. 3B**). These region clusters showed high signals in CRF pair Clusters 11 and 12 (**Fig. 3A**), which contains CRF pairs involving H3K27me3. Genes associated with these region clusters are mostly repressed and showed the lowest gene expression among all region clusters.

Using the ENCODE ChIP-seq data obtained from 309 TFs in K562 cells, we conducted a TF binding site enrichment analysis to investigate whether specific TFs preferentially bind to distinct genomic region clusters (**Fig. 3C, Supplementary Fig. S3B**). Our results indicated that region clusters with higher signals in CRF pairs associated with euchromatin marks (i.e., Clusters A-I) exhibited greater enrichment for most TFs and RNAPII reads compared to those with higher signals in CRF pairs related to heterochromatin marks (i.e., Clusters J-O).

The analysis also revealed cluster-specific TF binding patterns. For instance, Clusters A-F are enriched for binding sites of SP1, MAX, RNAPII, and several other TFs, consistent with their high expression activities and high percentages of PLS (**Fig. 3C**). These clusters also showed strong signals in CRF pair Clusters 1 and 2 (**Fig. 3A**), which primarily consist of H3K4me3-involved pairs. In Cluster G, in addition to a high enrichment of RNAPII binding sites, there is also notable enrichment for SFPQ, EWSR1, TAF15, HNRNPH1 and HNRNPUL1, transcription factors known for their functions in regulating transcription and RNA splicing [27–32] (**Fig. 3C**). Interestingly, Cluster G has elevated signals in CRF pair Clusters 4 and 5 (**Fig. 3A**), which mostly contain RNAPII-involved pairs. The enhancer-rich Clusters H and I are enriched in the binding sites of GATA1, GATA2, and STAT5A (**Fig. 3C**), which are TFs essential for blood cell development [33–35] and highly relevant to K562, a leukemia cell line.

The heterochromatin-related region clusters exhibited two major TF binding patterns. For instance, region Clusters J and K are significantly enriched for binding sites of SETDB1, an H3K9me3 writer, and CBX5, which binds to H3K9me3-marked regions, promoting chromatin compaction [36, 37] (**Fig. 3C**). This finding aligns with Clusters J and K and their unique pattern of high signals in CRF Clusters 10 (**Fig. 3A**), which primarily involves CRF pairs associated with H3K9me3. In contrast, the enriched binding sites in Clusters L, M, N and O, which are linked to CRF pairs involving H3K27me3, mostly belong to EZH2, MCM3, MCM5, CBX2, and CBX8 (**Fig. 3C**). This is consistent with the role of EZH2 as an H3K27me3 writer that facilitates heterochromatin formation. Both CBX2 and CBX8 are components of Polycomb complex and negatively regulate transcription [38]. Members of the mini-chromosome maintenance (MCM) protein family (e.g., MCM3, and MCM5) are involved in the initiation of eukaryotic genome replication at the replication origins, where the local transcription is often temporarily paused or reduced [39]. This analysis clearly reveals the physical association between a histone mark, its modification enzyme, and its reader.

### Comprehensive chromatin landscape obtained with Hi-Plex CUT&Tag reveals context-dependent functions of co-localized CRF pairs

To investigate how co-localized CRF pairs contribute to local transcriptional activity, we modeled the relationship between Hi-Plex CUT&Tag signals and gene expression levels through machine learning. Specifically, we extracted the Hi-Plex CUT&Tag signals of 501 CRF pairs at the promoter region (±1 kb of the transcription start site (TSS)) for each gene and used them to predict the expression levels of 19,718 protein-coding genes as measured by RNA-seq (**Methods**). We employed Random Forest (RF) [40], a non-parametric approach capable of modeling non-linear relationships, to build a prediction model, which achieves a high Pearson correlation between predicted and observed gene expression in cross-validation (*r* = 0.74; **Fig. 4A**).

**Figure 4.**
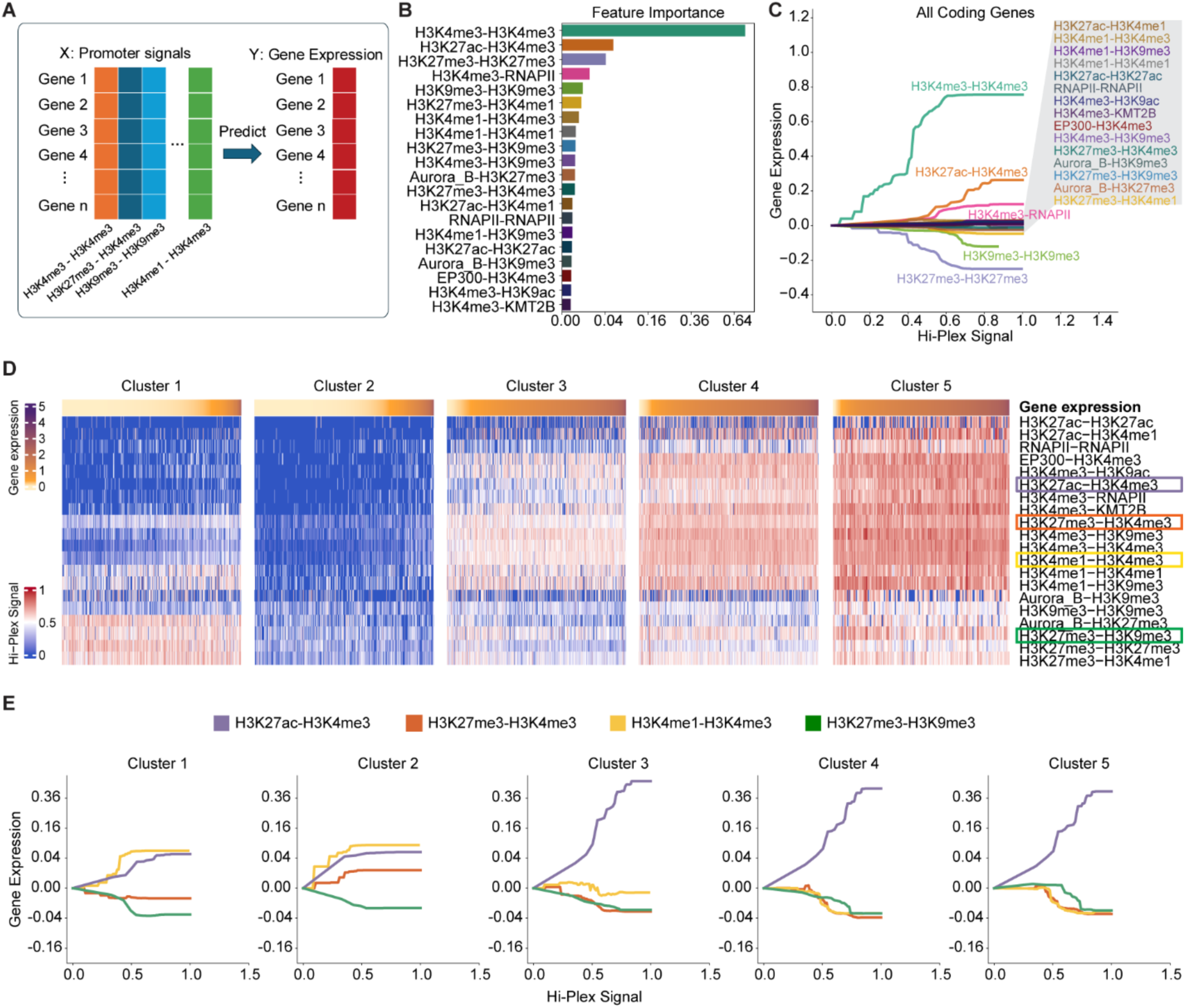
Machine learning model analysis reveals context-dependent relationships between CRF pairs and gene expression. **A.** Random Forest (RF) model trained and evaluated by five-folds cross-validation using CRF pair promoter signals as predictors (X) and gene expression as the outcome (Y). **B.** Bar plot of feature importance for the top 20 CRF pairs predicting gene expression. **C.** Partial dependence plots from the RF model illustrating the relationship between individual CRF pairs and gene expression. **D.** Heatmaps of five distinct clusters representing chromatin landscapes defined by all CRF pairs. Signals from the top 20 most important pairs are shown; clusters are ordered by average gene expression (low to high). **E**. Partial dependence plots showing relationships between four CRF pairs (H3K27ac–H3K4me3, H3K27me3–H3K4me3, H3K4me1–H3K4me3, and H3K27me3– H3K9me3) and gene expression across clusters.

Analysis of feature importance from the fitted RF model revealed that many of the top-ranked features are CRF pairs involving H3K4me3, consistent with its role in regulating gene expression as a mark of active promoters, and CRF pairs involving H3K27me3 and H3K9me3, consistent with their roles in repressing gene expression (**Fig. 4B**). Interestingly, many top-ranked CRF pairs are heterotones, including H3K27ac-H3K4me3, H3K4me3-RNAPII, H3K4me1-H3K4me3, as well as pairs of CRFs with opposing functions, such as H3K27me3-H3K4me1 and H3K4me3-H3K9me3. To further investigate the impact of these CRF pairs on gene expression (i.e., positive or negative effects), we examined how increasing the Hi-Plex CUT&Tag signal of each CRF pair in the fitted model influenced gene expression while holding all other features constant and visualized the relationship using partial dependence plots (**Fig. 4C**). We observed that CRF pairs composed of euchromatin marks, such as H3K4me3–H3K4me3, H3K27ac–H3K4me3, and H3K4me3–RNAPII, exerted the strongest positive effects on gene expression. In contrast, pairs involving heterochromatin marks, such as H3K27me3–H3K27me3 and H3K9me3–H3K9me3, were associated with the strongest negative effects.

Next, we investigated whether CRF pairs exhibit distinct regulatory patterns under different chromatin landscapes. To this end, we performed unsupervised clustering of genes based on promoter-associated Hi-Plex CUT&Tag signals across all CRF pairs. This analysis identified five gene clusters (Clusters 1–5) that reflected a progressive shift in chromatin state from closed to open. Correspondingly, gene expression levels increased gradually from Cluster 1 to Cluster 5 (**Fig. 4D**).

Cluster 1 was enriched for signals from heterochromatin-associated CRF pairs (e.g., H3K27me3-H3K9me3 and H3K27me3-H3K27me3) and depleted for signals from euchromatin-associated pairs (e.g., H3K4me3-H3K4me3 and H3K27ac-H3K4me3). Genes in this cluster exhibited low expression. Cluster 2 showed intermediate heterochromatin signals but low euchromatin signals, with genes also displaying low expression. Cluster 3 exhibited intermediate signals for both heterochromatin- and euchromatin-associated pairs and correspondingly moderate expression. Clusters 4 and 5 were characterized by high signals from euchromatin-associated pairs, with genes showing the highest expression levels.

Notably, while many CRF pairs showed consistent positive or negative correlations with gene expression across clusters, a subset displayed context-dependent relationships. For example, H3K4me1–H3K4me3 was associated with increased gene expression in Clusters 1 and 2, where euchromatin signals (e.g., H3K4me3, H3K4me1, and H3K27ac) were low, but with decreased expression in Clusters 3–5, where euchromatin signals were moderate to high (**Fig. 4E**). Similarly, the bivalent pair H3K27me3–H3K4me3 had little effect in Cluster 1, a weak positive effect in Cluster 2, and strong negative effects in Clusters 3–5 (**Fig. 4E**). By contrast, H3K27me3– H3K9me3 consistently correlated negatively with expression, whereas H3K27ac–H3K4me3 consistently correlated positively. Other pairs, such as H3K4me3–H3K4me3, H3K4me3–RNAPII, and H3K27ac–H3K27ac, showed consistent positive associations, while H3K27me3–H3K27me3 and H3K9me3–H3K9me3 showed consistent negative associations. Still others, including H3K4me3–H3K9me3 and H3K4me1–H3K9me3, exhibited context-dependent effects (**Supplementary Fig. S4**).

Together, these results suggest that, rather than functioning solely as positive or negative regulators, many co-localized CRF pairs influence gene expression in a context-dependent manner, shaped by the chromatin landscapes captured by Hi-Plex CUT&Tag.

### Context-dependent gene regulation model enhances prediction of perturbation-induced gene expression changes

We hypothesized that if the context-dependent gene regulation model holds true, it could enable more accurate predictions of gene expression changes following perturbations to the chromatin landscape. To test this hypothesis, we treated K562 cells with sodium butyrate, a pan-histone deacetylase (HDAC) inhibitor, known to induce global chromatin remodeling [41]. We then performed Hi-Plex CUT&Tag using the same cocktail of 36 barcoded mAbs and RNA-seq on both sodium butyrate-treated and untreated K562 cells (**Fig. 5A**).

**Figure 5.**
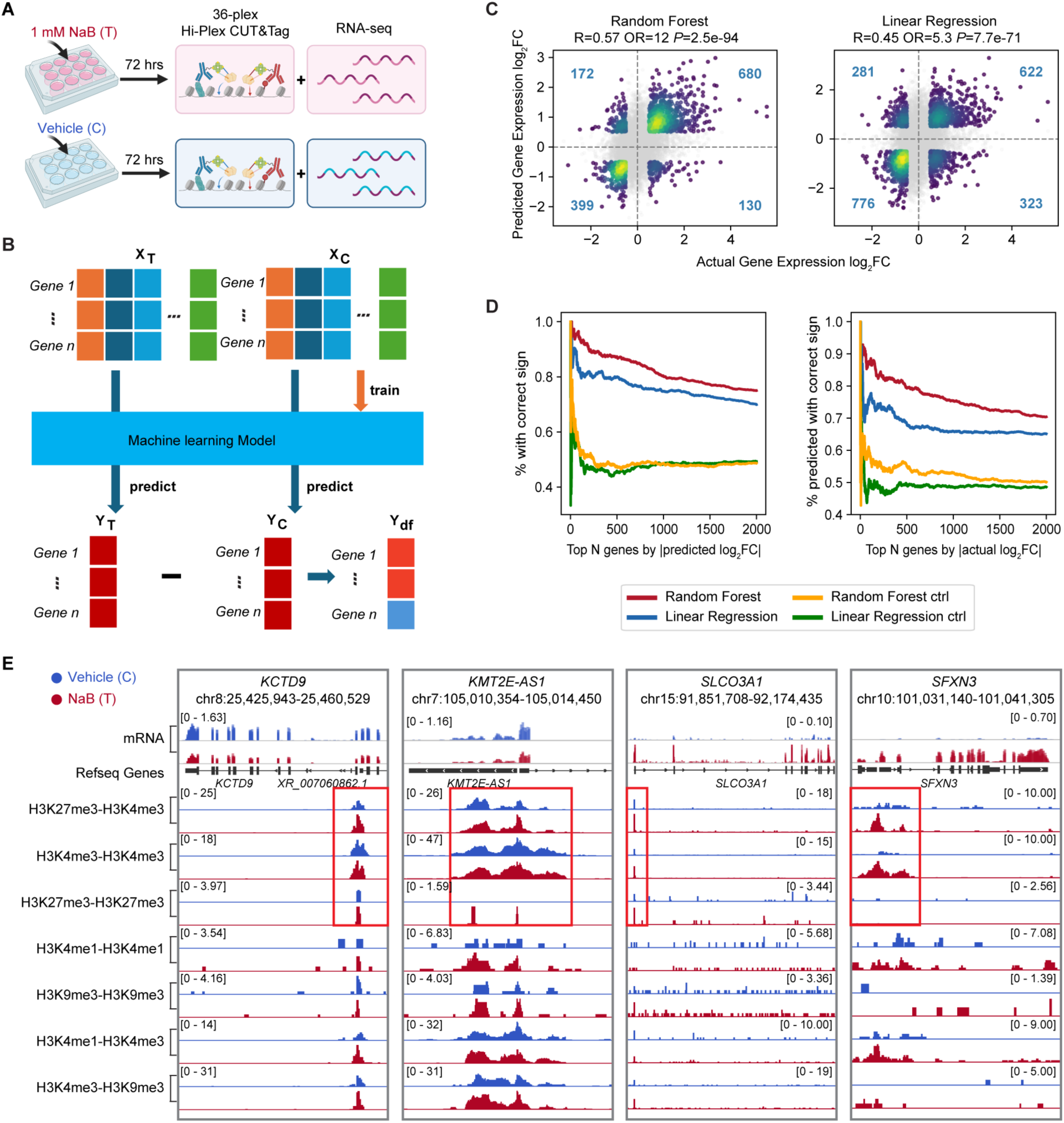
Dynamic changes in Hi-Plex CUT&Tag predict gene expression changes between different conditions. **A.** Global chromatin remodeling induced with sodium butyrate treatment in K562 cells. Hi-Plex CUT&Tag and RNA-seq were performed in parallel on both treated (T) and untreated (C) samples. **B.** A Random Forest (RF) model is trained using Hi-Plex signals from untreated samples to predict gene expression. The trained model is then applied to predict gene expression changes between treated and untreated samples based on Hi-Plex signals. **C.** Scatterplots comparing the true and predicted log_2_ fold change of gene expression between treated and untreated samples for Random Forest and Linear Regression, respectively. **D.** Left: Percentage of top-N differential genes (based on predicted log2 fold change) whose directions were correctly predicted by Random Forest, Linear Regression, and control models. Right: Percentage of top-N differential genes ranked by RNA-seq log2 fold change with correctly predicted directions. **E.** Signal tracks showing the gene expression and Hi-Plex CUT&Tag signals near genes *KCTD9*, *KMT2E-AS1*, *SLCO3A1*, and *SFXN3* before (blue) and after (red) treatment.

First, we validated the presence of dynamic changes in the chromatin landscape before and after treatment using Hi-Plex CUT&Tag. Consistent with the known effects of sodium butyrate, the DNA fragment length distribution shifted towards shorter fragments (**Supplementary Fig. S5B, Supplementary Note**). Differential analysis identified significant changes in 14 of the 16 CRF pair clusters (**Supplementary Fig. S5C-E; Supplementary Note**).

Next, we trained both the random forest (RF) model and a multiple linear regression model using Hi-Plex CUT&Tag and RNA-seq data from untreated cells. The trained models were then applied to the post-treatment Hi-Plex CUT&Tag data to predict gene expression changes, with prediction accuracy assessed by comparing to post-treatment RNA-seq data (**Fig. 5B**). In the linear model, the effect of each CRF pair on gene expression is a constant characterized by the corresponding regression coefficient, and thus, it is context-independent. We also trained random prediction models as controls (**Methods**).

Among these models, the RF model consistently outperformed both the linear and random control models. It achieved higher Pearson correlation coefficients between predicted and true log_2_ gene expression fold changes (log_2_FC) (r = 0.57 for RF vs. 0.45 for linear model), greater odds ratios (OR) for predicting the correct direction of change (OR = 12 for RF vs. 5.3 for linear model), and superior precision and recall (**Fig. 5C, D; Supplementary Fig. S6B**). The superior performance of the RF model indicates that context-independent constants in the linear model are insufficient to capture the effects of CRF pairs on gene expression, validating the presence of non-linear context-dependent relationships between them.

The complex interplay among CRF pairs and their context-dependent effects on gene expression highlights the importance of comprehensive epigenomic profiling in understanding CRF functions. For instance, the bivalent CRF pair H3K27me3-H3K4me3 showed increased signal intensity at the promoters of four genes, *KCTD9*, *KMT2E-AS1*, *SLCO3A1*, and *SFXN3*, following sodium butyrate treatment (**Fig. 5E**). Despite this shared chromatin signature, *KCTD9* and *KMT2E-AS1* were transcriptionally downregulated, while *SLCO3A1* and *SFXN3* were upregulated after treatment (**Fig. 5E**). While H3K27me3-H3K4me3 alone cannot explain these discrepancies,

examining additional CRF pairs revealed distinct chromatin contexts for these genes. For example, H3K4me3 homotone signals increased at the *SLCO3A1* and *SFXN3* promoters after treatment but remained unchanged at *KCTD9* and *KMT2E-AS1* promoters. In contrast, H3K27me3 homotone signals increased at *KCTD9*, *KMT2E-AS1*, and *SLCO3A1* after treatment but remained unchanged at *SFXN3* (**Fig. 5E**). Other CRF pairs (e.g., H3K4me1-H3K4me1, H3K9me3-H3K9me3, H3K4me1-H3K4me3, H3K4me3-H3K9me3) exhibited similarly complex and non-uniform dynamics (**Fig. 5E**). These examples underscore the multifaceted nature of chromatin regulation and demonstrate that changes in individual chromatin marks or pairs of marks cannot reliably predict gene expression outcomes without considering the broader epigenetic context. This context likely involves numerous CRFs and their co-localization patterns. While Hi-Plex CUT&Tag can profile this multidimensional landscape in a single assay, such comprehensive profiling would be difficult using traditional single-protein or low-plex technologies.

### Hi-Plex CUT&Tag profiling in single cells reveals synergy across co-localized CRF pairs

Previous studies have demonstrated the utility of CUT&Tag and multi-CUT&Tag for profiling chromatin regulators at the single-cell level [6–8, 15, 16]. Building on these approaches, we have developed protocols to adapt Hi-Plex CUT&Tag for single-cell profiling. We pooled a panel of 16 barcoded mAbs, targeting six histone PTMs (H3K4me3, H3K9me3, H3K9ac, H3K14ac, H3K27me3, and H3K27ac) and 10 TFs (CTCF, RNAPII S2P, c-Jun, c-Fos, Max, Myc, USF1, USF2, NRF1, and YY1). Again, rabbit IgG was included as a negative control. To generate single-cell Hi-Plex CUT&Tag data in K562 cells, we performed Hi-Plex CUT&Tag assay in bulk until the tagmentation phase was completed. Next, we sorted single nucleus into 384-well plates and prepared single-cell libraries in each well with distinct index primer pairs. The pooled single-cell libraries were then sequenced. Two replicates were performed, consisting of 1,278 and 1,318 cells, respectively, after quality control (**Methods**).

To evaluate data quality, we pooled single-cell data into pseudobulk data and found that the pseudobulk data closely matched the corresponding bulk Hi-Plex CUT&Tag data from the same CRF pairs (**Fig. 6A, B**). For example, the correlation between bulk and pseudobulk samples was 0.93, 0.95, and 0.62 for H3K4me3-H3K4me3, H3K27me3-H3K27me3, and H3K27me3-H3K4me3, respectively (**Fig. 6B**). Additionally, combinatorial co-occurrence patterns of CRF pairs were highly similar between bulk and pseudobulk data (**Fig. 6C**). This consistency demonstrates the high quality of the single-cell data.

**Figure 6.**
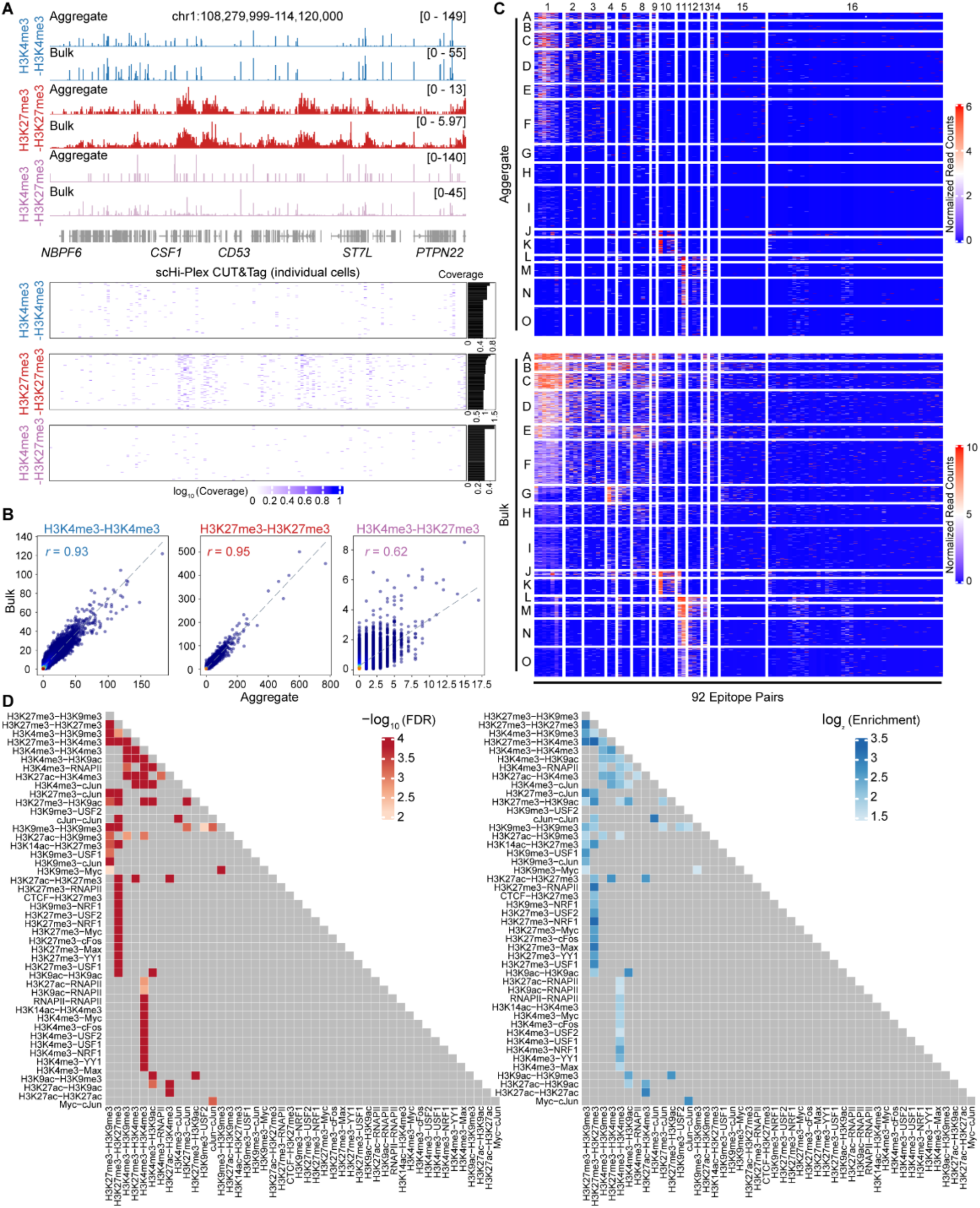
Single-cell Hi-Plex CUT&Tag reveals co-occurrence of CRF pairs in the same cell. **A.** Signal track comparison between aggregated single-cell and bulk Hi-Plex CUT&Tag data for two homotone CRF pairs (H3K4me3-H3K4me3 and H3K27me3-H3K27me3) and one heterotone pair (H3K4me3-H3K27me3). The lower panel displays log_10_-transformed read coverage from the top 100 single cells (ranked by total read count) for each corresponding CRF pair. **B.** Scatter plots showing strong genome-wide correlations between aggregated single-cell and bulk data for H3K4me3-H3K4me3, H3K27me3-H3K27me3, and H3K4me3-H3K27me3, with Pearson’s correlation coefficients of 0.93, 0.95, and 0.62, respectively. **C.** Aggregated single-cell data recapitulate the combinatorial co-localization patterns observed in bulk. The upper panel shows aggregated single-cell signals; the lower panel shows bulk signals across the same CRF pairs and region clusters defined in bulk data analysis (as in Fig. 3A). **D.** Co-occurrence analysis of CRF pair combinations across genomic regions used in the bulk analysis (i.e., regions in Fig. 3A), considering only CRF pair combinations with > 5 observed co-occurrence events. A total of 46 significant co-occurring CRF pair combinations were identified (FDR < 0.05, enrichment score > 1.5). The left panel shows –log_10_-transformed FDR values; the right panel shows log_2_-transformed enrichment scores.

Although bulk Hi-Plex CUT&Tag analysis identifies co-occurrence patterns of multiple CRF pairs within the same genomic regions (**Fig. 3A**), these data cannot distinguish between the following two scenarios: (1) the multiple CRF pairs have synergistic functions, requiring co-occurrence in the same cell, or (2) the CRF pairs have independent functions, occurring in distinct cell subpopulations but appearing co-localized in pooled bulk data.

Single-cell Hi-Plex CUT&Tag resolves this ambiguity by enabling direct evaluation of these scenarios. For instance, heterotone pairs H3K4me3-H3K9ac and H3K4me3-H3K9me3, both from CRF pair Cluster 1, frequently co-occur in the same genomic regions in bulk analysis (**Fig. 3A**). However, it remains unclear whether these CRF pairs occur in mutually exclusive sets of cells or co-occur within the same cells. Using single-cell data, we assessed their cell-level co-occurrence by counting events where both CRF pairs were observed within the same genomic regions and in the same cells. These observed counts were compared to random expectations based on a permutation-derived null distribution (**Methods**). The analysis revealed significant co-occurrence of CRF pairs in the same regions and cells (**Fig. 6D**), indicating their potential synergistic roles in regulating gene expression. Applying this approach to all paired combinations of CRF pairs whose observed co-occurrence events are larger than 5, we found that 46 pairs of them showed significant co-occurrence (**Fig. 6D**, FDR < 0.05, enrichment score > 1.5). Notable examples of significant co-occurrence pairs include: (1) H3K27me3-H3K9me3 and H3K4me3-H3K9me3; (2) H3K4me3-H3K4me3 and H3K4me3 heterotones such as H3K4me3-H3K9ac, H3K4me3-RNAPII, H3K4me3-cJun, and H3K14ac-H3K4me3; (3) H3K27me3-H3K27me3 and H3K27me3 heterotones, including H3K27me3-H3K4me3, H3K27me3-cJun, H3K27me3-H3K9ac, H3K14ac-H3K27me3, H3K27me3-RNAPII, H3K27me3-MYC, H3K27me3-MAX, and H3K27me3-USF1.

These results highlight the capability of Hi-Plex CUT&Tag to analyze the chromatin landscape at single-cell resolution.

## DISCUSSION

It has become increasingly clear that a key to understanding a global picture of how nuclear proteins and histone PTMs synergize to guide chromatin function is the ability to interrogate cross-talk among these factors while concurrently observing their detailed positional information across the genome. Querying cross-talk on chromatin has been technically challenging until now. Here, we demonstrate the utility for Hi-Plex CUT&Tag, showing that this highly multiplexed, genome-wide sequencing approach enables the simultaneous detection of hundreds of chromatin-associated interactions occurring on the same chromatin. Our ability to multiplex 36 barcoded mAbs relies on minimizing background noise and cross-contamination by barcoding the mAbs using the nearly irreversible biotin-streptavidin interaction. Additionally, we optimized the stoichiometry of biotinylation, ensuring that most antibodies had fewer than two biotin moieties per antibody. This adjustment enhanced the likelihood of detecting co-localization events (i.e., heterotone reads). Notably, over 68% of the reads were heterotone, reflecting a high proportion of unambiguous co-localization events. Accordingly, Hi-Plex CUT&Tag represents a novel technology that advances multiplex chromatin profiling with enhanced specificity and sensitivity. The approach enables simultaneous analysis of co-localized chromatin events across hundreds of CRF pairs. Each sequenced DNA fragment produced by this method encodes information about the identities of two CRFs co-localized to the same strand of DNA, their genomic locations, and the molecular distance between them.

This rich dataset facilitates a comprehensive characterization of the chromatin landscape in a context-dependent manner. We believe this technology could also be used in the future to establish causal relationships between dynamic changes of histone marks and recruitment of histone modifiers and/or TFs, in connection to transcriptional outcome, with temporal resolution. For example, during iPSC differentiation into neural progenitor cells, this technology could be used to uncover causal relationships between the dynamic loss of H3K27me3 at lineage-specific gene promoters and the recruitment of demethylases such as KDM6A/B, along with transcription factors like SOX2 or PAX6, thereby linking specific epigenetic remodeling events to transcriptional activation with temporal resolution.

Using this technology, we have, for the first time in the same sample, systematically mapped co-localized events across more than 500 CRF pairs, including numerous heterotone pairs and bivalent marks. We characterized the properties of these CRF pairs, such as associations with *cis*-regulatory elements, repetitive sequences, DNA methylation, and gene expression. Notably, we identified distinct TF binding patterns associated with different combinatorial chromatin co-localization states, suggesting that specific chromatin configurations correspond to distinct regulatory programs.

This ability to comprehensively map chromatin landscapes enables a systematic exploration of the relationships between co-localized eCRF pairs and gene expression. Our analyses revealed a complex, context-dependent relationship between co-localized CRF pairs and gene expression. For example, the co-localization of H3K4me3 and H3K4me1 on average is positively associated with gene expression in an overall repressive epigenetic context with low gene expression but shows negative association with gene expression in an active epigenetic context with high gene expression. One possible explanation is that the H3K4me3-H3K4me1 pair marks a transition from a repressive state to a poised-to-activation state (positively correlating with gene expression when expression is low), or a transition from an active state to a poised-to-silence state (negatively correlating with gene expression when expression is high). On the other hand, the vast majority of co-localized CRF pairs did not significantly contribute to transcriptional regulation (**Fig. 4**), suggesting that not all histone marks or chromatin-associated proteins have equal regulatory impact. However, we cannot rule out the possibility that certain CRF pairs may play critical roles in transcriptional regulation in other cellular contexts or in response to different perturbations or stimuli, as our analysis was limited to a single cell line. To further elucidate the functional roles of co-localized histone marks and chromatin-associated proteins, future studies should expand to include multiple cell types and tissues, and a broader range of perturbations or environmental stimuli. Integrating temporal profiling with high-resolution epigenomic and transcriptomic analyses will be essential to capture dynamic regulatory events. Additionally, functional validation using genetic or chemical perturbation of candidate CRF pairs will help determine their causal impact on transcriptional outcomes.

Our findings highlight the limitations of traditional approaches that analyze individual histone modifications in isolation. Simplistic interpretation, such as “histone mark A is positively correlated with gene expression” or “histone mark B is repressive,” overlook the nuanced, combinatorial interactions among chromatin marks and chromatin-associated factors that govern transcriptional regulation. A precise functional understanding of a given CRF requires consideration of its broader chromatin context, including the presence or absence of neighboring histone modifications, histone remodelers, and/or TFs. Hi-Plex CUT&Tag is uniquely positioned to address this challenge. By simultaneously profiling hundreds of CRF pairs, it enables a comprehensive analysis of chromatin context. Moreover, its capacity to measure co-localized histone marks makes it particularly powerful for investigating the functional roles of CRF pairs, including bivalent modifications, which are extremely challenging to assess using conventional single-mark profiling methods.

Furthermore, we have developed single-cell Hi-Plex CUT&Tag, which enables the analysis of synergy between co-localized CRF pairs by examining their co-occurrence within individual cells. As with other single cell technologies, current data from individual cells are sparse. However, we anticipate that continued optimization of experimental protocols, along with advancements in computational algorithms, will significantly enhance data quality. As a result, single-cell Hi-Plex CUT&Tag is poised to become a powerful tool for high-resolution, comprehensive studies of chromatin landscapes. We envision that by coupling scRNA-seq with single-cell Hi-Plex CUT&Tag in the same cell will enable integrated, multi-dimensional profiling of chromatin states and gene expression, ultimately providing deeper insights into the regulatory mechanisms driving cell fate decisions and cellular heterogeneity.

## Supporting information

Hi-Plex CUT&Tag Supplementary materials

## ACKNOWLEDGEMENTS

This work was supported in part by NIH grant 5R01HL149961-04, 5R01NS127913, 1P50 AA027054, R01EY030475, R24AA025017, R01EY035381 to HZ; R01HG013409 to HJ;R01GM118760, R01CA236209, P20GM121293, and funds from the Winthrop P. Rockefeller Cancer Institute and the Arkansas Biosciences Institute to SDT. We are grateful to CDI Labs and PTM Bio for their technical support.

## MATERIAL & METHODS

### Biological materials & reagents

K562 cells were cultured in RPMI medium (Gibco, 11875119), supplemented with 10% FBS (Gemini Bio, 100-602-500) and 1% penicillin-streptomycin (ThermoFisher, 15140122). For sodium butyrate treatment, actively growing K562 cells were seeded in six-well plates at a density of 0.1 × 10⁶ cells/mL. Sodium butyrate (Millipore Sigma, 19-137) was added to a final concentration of 1 mM, and cells were incubated for 72 hours. Distilled water (ThermoFisher, 10977023) was added to control cells as a vehicle control. All antibodies used in this study are listed in **Supplementary Table S3**, and all reagents and materials are listed in **Supplementary Table S4**. DNA oligonucleotides for barcoding and sequencing were designed based on previously published sequences [14, 15], ordered from Integrated DNA Technologies, and are listed in **Supplementary Tables S5** and **S6**.

### Antibody barcoding

Each antibody was resuspended in PBS to a final concentration of 1 mg/mL and incubated with NHS-PEG12-Biotin (ThermoFisher, A35389) at a molar ratio of 1:5 overnight at 4 °C. The following day, unreacted NHS-PEG12-Biotin was removed using Zeba™ Spin Desalting Columns or plates (40K MWCO, ThermoFisher, 87767, 87775). Biotinylated antibodies were barcoded with a mixture of P5 and P7 adaptor sequences. Each adaptor oligonucleotide contained an Illumina read 1 (P7 adaptor) or read 2 (P5 adaptor) primer sequence, an 11-nucleotide DNA barcode sequence, and a double-stranded Tn5 binding mosaic end. The adaptor sequences are listed in **Supplementary Table S5**. The P5 adaptor oligonucleotides consisted of the “11nt_Bio-P5-N” series, and the P7 adaptor oligonucleotides consisted of the “11nt_Bio-P7-N” series, where “N” represents a specific barcode identifier. P5 or P7 adaptor oligonucleotides (50 µL of 100 µM) were annealed with Tn5MErev oligonucleotides (10 µL of 500 µM) in 40 µL of distilled water (ThermoFisher, 10977023) by incubating at 95 °C for 2 min, followed by slow cooling to room temperature. Equal amounts of P5 and P7 adaptors were then mixed to generate the adaptor mixture. To barcode each antibody, the biotinylated antibody was diluted 10-fold in PBS, then streptavidin (ThermoFisher, 21122) and the adaptor mixture were added at a molar ratio of 1:1:2 (antibody:streptavidin:adaptor), and the reaction was incubated at room temperature for 1 hour. To saturate binding sites on streptavidin, D-biotin (ThermoFisher, B20656) was added to a final concentration of 2.25 µM, and the mixture was incubated at room temperature for 30 min. All barcoded mAbs were then pooled and concentrated using 30K Amicon centrifugal filters (Millipore Sigma, UFC503096, UFC803096). The mAb mixture was stored at 4 °C until use.

All 11-nucleotide DNA barcode sequences were initially designed using our established algorithm with a Hamming distance of 3 [42]. Here, we further filtered them and retained only barcodes with a Hamming distance greater than 5. The retained barcodes allowed for up to 2 nucleotide mismatches during demultiplexing. We also performed primer dimer analysis using the Multiple Primer Analyzer [43] to exclude barcodes that would anneal with other barcodes or themselves at room temperature.

### Hi-Plex CUT&Tag

Concanavalin A beads were first activated by washing twice in binding buffer (20 mM HEPES pH 7.5, 10 mM KCl, 1 mM CaCl₂, 1 mM MnCl₂). Actively growing K562 cells (100,000 cells) were washed once in PBS and once in wash buffer (20 mM HEPES pH 7.5, 150 mM NaCl, 0.5 mM spermidine, 1× protease inhibitor cocktail) and then resuspended in 0.5 mL of wash buffer and transferred to 10 µL of activated Concanavalin A beads. Cells were incubated with the beads at room temperature for 15 min on a rotator. The buffer was removed using a magnetic stand. Cell-bound beads were resuspended in 100 µL of wash buffer with the antibody mixture (1 µg of each antibody), 0.05% digitonin, and 2 mM EDTA, and then incubated at room temperature for 1 hour or at 4 °C overnight on a rotator. The beads were washed four times in Dig-med buffer (20 mM HEPES at pH 7.5 with 300 mM NaCl, 0.5 mM spermidine, 0.01% digitonin, and 1× protease inhibitor cocktail), then resuspended in 100 µL of Dig-med buffer containing 10 mM MgCl₂ and 5 µg of Tn5 (Diagenode, C01070010-20), and incubated at 37 °C for 1 hour on a rotator. Tagmentation was stopped by adding 3.33 µL of 0.5 M EDTA, 1 µL of 10% SDS, and 0.33 µL of 20 mg/mL Proteinase K (ThermoFisher, EO0491). The sample was then vortexed and incubated at 50 °C for 1 hour. DNA was purified by phenol-chloroform extraction. Phenol-chloroform-isoamyl alcohol (100 µL, pH 8, ThermoFisher, 17908) was added, mixed thoroughly, transferred to phase-lock tubes (ThermoFisher, NC1093153), and centrifuged for 3 min at room temperature at 16,000 × *g*. Chloroform (100 µL) was added to the aqueous phase, mixed, and centrifuged for 5 min at 16,000 × *g*. The aqueous phase was transferred to a new tube and precipitated with 250 µL of 100% ethanol and 8.75 µL of 20 mg/mL glycogen at −80 °C overnight. The following day, samples were centrifuged for 15 min at 4 °C at 16,000 × *g*. The DNA pellet was washed with 1 mL of 100% ethanol, centrifuged for 5 min at 4 °C at 16,000 × *g*, air-dried, and dissolved in 23 µL of 10 mM Tris-HCl (pH 8) containing 1% (v/v) RNase A (ThermoFisher, EN0531). The sample was incubated for 10 min at 37 °C. For library amplification, purified DNA (21 µL) was mixed with barcoded i5 and i7 primers (2 µL each at 10 µM) and NEBNext Ultra II Q5 Master Mix (25 µL, NEB, M0544S). The mixture was amplified using the following thermocycler program: 72 °C for 5 min, 98 °C for 30 sec; 17 cycles of 98 °C for 10 sec, 63 °C for 10 sec; 72 °C for 1 min; hold at 4 °C. Libraries were purified using AMPure XP beads (Beckman, A63881) at a 1:1.1 sample-to-bead ratio according to the manufacturer’s protocol.

i5 primer:

5’-AATGATACGGCGACCACCGAGATCTACACNNNNNNNNTCGTCGGCAGCGTC-3’

(N: 8-nucleotide barcode) i7 primer:

5’-CAAGCAGAAGACGGCATACGAGATNNNNNNNNGTCTCGTGGGCTCGG-3’

(N: 8-nucleotide barcode)

### Single-cell Hi-Plex CUT&Tag protocol

To set up single-cell Hi-Plex CUT&Tag assays, 100,000 K562 cells were collected and washed once with PBS and once with wash buffer. Cells were resuspended in 1 mL of NP-wash-buffer (wash buffer with 0.01% Digitonin, 0.01% NP-40, and 20 mM sodium butyrate) and incubated with activated Con-A beads for 15 min at room temperature on a rotator. After removing the buffer, beads were resuspended in 100 µL of NP-wash-buffer with 2 mM EDTA, incubated with antibody mixture for 1 h at room temperature, and washed four times with NP-Dig-med buffer (Dig-med buffer with 0.01% NP-40). Beads were then resuspended in 100 µL of NP-Dig-med buffer containing 10 mM MgCl₂ and 5 µg of Tn5 and incubated at 37 °C for 1 h.

Following tagmentation, cell-bead complexes were resuspended in 1 mL of 10 mM Tris-HCl containing 10 µg/mL DAPI and filtered through a cell strainer into round-bottom tubes. Single cells were sorted into 384-well plates using a MoFlo XDP sorter, centrifuged for 3 min at 3000 × *g* and 4 °C, and stored at −80 °C. For processing, 1 µL of 0.095% SDS was added to each well using an Echo 650 Acoustic Liquid Handler, followed by centrifugation and incubation at 58 °C for 1 h. Next, 0.5 µL of 2.5% Triton X-100 and 0.5 µL i5/i7 primer mix (10 µM) were added to each well, providing unique index pairs. After adding 2 µL of NEBNext Ultra II Q5 Master Mix (NEB, M0544S), each single cell library was amplified using the following program: 58 °C for 5 min, 72 °C for 5 min, 98 °C for 30 sec; 17 cycles of 98 °C for 10 sec, 63 °C for 10 sec; 72 °C for 1 min; hold at 4 °C.

Single cell libraries were then pooled using Single-well Deep Well Plates (Miltenyi Biotec, 130-114-966). Each 384-well plate was inverted over the deep-well plate, centrifuged for 1 min at 1000 × *g* and 4 °C, and the pooled library was transferred to a new tube. Cleanup was performed with AMPure XP beads (Beckman, A63881) at a 1:1.1 sample-to-bead ratio according to the manufacturer’s instructions.

### RNA-seq library preparation

Total RNA was purified using the miRNeasy Mini Kit (Qiagen, 217004) following the manufacturer’s instructions for animal cells. mRNA was then purified using the Dynabeads™ mRNA Purification Kit (Invitrogen, 61006). mRNA sequencing libraries were prepared using the NEBNext® Ultra™ II RNA Library Prep Kit for Illumina (NEB, E7770S) following Section 1 instructions. Libraries were indexed using NEBNext® Multiplex Oligos for Illumina® (Index Primers Set 1) (NEB, E7335S).

### Data demultiplexing and alignment

After the NextGen sequencing, the paired-end sequencing reads were demultiplexed into different chromatin regulatory factor (CRF) pairs based on the corresponding barcodes from the two ends of each read using CutAdapt [44] with the following parameters: The maximum number of allowed errors was set to 2 (-e 2). To ensure stringent matching, indels were not allowed (--no-indels). The undetermined barcodes that did not match any of the designed CRFs were assigned with an “unknown” tag. After demultiplexing, heterotones, CRF pairs with two different barcode sequences (e.g., H3K27me3-H3K4me3 vs. H3K4me3-H3K27me3), were merged since they are biologically interchangeable. Therefore, for an experiment with *n* barcoded mAbs, there will be *n* homotones (i.e., CRF pairs with the same barcode sequences) and *n* * (*n* - 1) / 2 heterotones. The demultiplexed reads were then aligned to the human reference genome hg38 using Bowtie2 [45, 46] with the following parameters: the minimum and maximum fragment lengths were set to 10 and 800 respectively (-I 10 -X 800). The first 45 bases from the 5’ end of each read (−5 45) were trimmed to remove barcode and adaptor sequences. The alignments were performed using the local alignment mode (--local), with the sensitivity preset set to very sensitive (--very-sensitive-local). Additionally, we excluded unaligned reads (--no-unal), mixed alignments (--no-mixed), and discordant alignments (--no-discordant).

### Quality control

After alignment, only the CRF pairs with enough reads were retained for further analyses. CRF pairs with total read counts lower than the first quantile of all CRF pairs were removed. As a result, 501 CRF pairs were retained where each CRF pair has at least 869 reads.

### Fragment type decomposition

The fragment length distribution for all CRF pairs exhibits three distinct modes: sub-nucleosome, mono-nucleosome, and di-nucleosome or more (di+-nucleosome). To define the range of these modes, a density curve was fitted to the distribution using the default settings of the hist function in R. The boundaries between the three fragment types were determined by identifying the local minima between adjacent peaks in the distribution density curve. Fragments shorter than 121 bp were classified as sub-nucleosome, fragments between 122 bp and 298 bp were classified as mono-nucleosome, and fragments longer than 299 bp were classified as di-nucleosome and larger (di+-nucleosome).

### Biclustering heatmap

To analyze combinatorial co-occurrence patterns of various CRF pairs within the same genomic regions, we performed a biclustering analysis of CRF pair signals across the genome. The genome was divided into 800 bp non-overlapping regions, and the number of reads in each region for each CRF pair was counted, resulting in a region-by-CRF-pair count matrix. To ensure comparability, the data were normalized and transformed as follows. Each CRF pair was normalized by its library size (i.e., total counts) and scaled by a factor of one million. The normalized counts were log_2_-transformed after adding a pseudocount of one to avoid log-zero. Quantile normalization was further applied to standardize the distribution across all CRF pairs.

Performing biclustering on genomic regions across the whole genome would introduce noise due to regions with extremely low signals. To reduce the noise, we first selected highly variable genomic regions and performed biclustering on these highly variable genomic regions. We then assigned the remaining genomic regions to the genomic region clusters obtained above. To select highly variable regions, a generalized additive model (GAM) curve was fitted to the log_2_-transformed mean and variance of each genomic region across all CRF pairs. Highly variable regions were selected based on two criteria: (1) Regions with average values in the top 1% were retained; (2) Regions were stratified into 100 strata based on a uniform split of the range of log_2_-transformed mean values, and the regions with variance in the top 1% in each stratum and above the GAM curve were retained. This process resulted in the selection of 8,536 genomic regions with the strongest signals and highest variability across 501 CRF pairs for biclustering analysis. Using the ComplexHeatmap R package [47], we applied k-means clustering, grouping the genomic regions into 15 clusters and the CRF pairs into 16 clusters. The combinatorial patterns of these highly variable genomic regions are shown in **Fig. 3A**, where rows are genomic regions and columns are CRF pairs.

Taking these 15 row clusters of highly variable genomic regions as seeds, we expanded our analysis to all other genomic regions. Among these, 3,463,883 regions had 10 or fewer CRF pairs with non-zero raw reads; we designated these as “background” regions and excluded them from further analyses. For the remaining regions, we first derived the mean signal profile for each row cluster and then computed the Pearson correlation between each non-background region and these profiles. Each region *i* was then associated with its most correlated cluster *c*, and the corresponding correlation *r_ic_*, was used to characterize the similarity between the region and the cluster. Using this procedure, all non-background regions were assigned to a cluster. However, some regions may show low correlation with their corresponding cluster. To ensure high-quality cluster assignment, we retained only regions with *r_ic_* values in the top 25% of the distribution of *r_ic_* across all non-background regions. This approach added 96,983 genomic regions to the seed clusters shown in **Fig. 3A**. Together with the original 8,536 seed regions, this yielded a total of 105,519 genomic regions, whose biclustering heatmap is shown in **Supplementary Fig. S3A**.

### Peak calling and genomic region annotation

To identify signal regions for each CRF pair, we performed peak calling using SEACR version 1.4 with its default setting [48]. To annotate the identified peaks and 105,519 genome-wide highly variable regions identified in **Supplementary Fig. S3A**, the following procedure was applied. Exons, introns, and untranslated regions (UTRs) were annotated using the annotatePeak function in ChIPseeker [49]. Cis-regulatory elements (cCREs) were annotated using the findOverlaps function in GenomicRanges R package [50] based on the ENCODE candidate Cis-Regulatory Elements database (cCRE) [21]. Repetitive elements were annotated using RepeatMasker [22]. Chromatin state annotations were obtained using ChromHMM [26].

### Prediction model

To investigate the relationship between CRF pair activities and gene expression, we trained a Random Forest (RF) model to predict gene expression (log_10_TPM) based on the activity of 501 CRF pairs in the promoter regions of corresponding genes. Outlier genes with expression above the 99.5^th^ percentile of all genes and genes with no detectable expression in the untreated condition (C) were excluded. The RF model was trained and evaluated using a 5-fold cross-validation procedure implemented with the scikit-learn package [51]. CRF pair activities were summarized as promoter activity scores by counting the number of reads within ±1 kb of the transcription start site (TSS) of each gene.

To evaluate the contribution of each CRF pair (i.e., feature) to gene expression prediction, feature importance scores were calculated using the feature_importances_attribute of scikit-learn’s RandomForestRegressor [51], which implements the mean decrease in impurity method. This method quantifies how much each CRF pair contributes to decreasing node impurity across all trees in the forest. For each split in every tree, the algorithm calculates the reduction in variance. The importance of each epigenomic feature is computed as the sum of these variance reductions across all nodes where that feature is used for splitting, normalized by the total number of trees. This approach provides a measure of each CRF pair’s contribution to explaining gene expression variability across the genome.

### Feature analysis using partial dependence plot

Partial dependence plots (PDPs) were generated to analyze and visualize the relationship between each CRF pair and gene expression within the fitted RF models. PDPs assess the effect of a specific CRF pair on the model’s predicted gene expression while holding all other CRF pairs constant. For a CRF pair of interest, its values were systematically varied across their observed range for each gene. The predicted gene expression values were then averaged across all genes to capture the overall impact of the CRF pair on gene expression. This approach provides an interpretable visualization of how changes in the activity of a single CRF pair influence gene expression, conditional on the epigenetic context defined by all other CRF pairs.

To further explore the context-dependent effects of each individual CRF pair on gene expression, we performed k-means clustering on genes based on all CRF pairs. The optimal number of clusters was determined using the elbow method implemented in Python kneed library [52]. For each gene cluster, partial dependence plots are calculated using the genes within that cluster.

### Differential expression prediction

The prediction models trained using data from the untreated condition (C) were applied to predict log_10_TPM of gene expression levels in both treated (T) and untreated (C) conditions. Predicted log_2_(fold changes) were calculated as: log_2_FC = (pred_y_T / log_10_(2)) - (pred_y_C / log_10_(2)), where pred_y_T and pred_y_C represent predicted expression levels in treated and untreated conditions, respectively. Log_2_FC prediction performance was evaluated using 5-fold cross-validation with the scikit-learn package [51]. Specifically, for each fold, the data were partitioned into a training set with 80% of genomic regions and a test set with the remaining 20% regions. Four models, namely Linear Regression, Random Forest, Linear Regression control, and Random Forest control, were trained on the training set and applied to the test set to make predictions. Their prediction performances were then compared. Control models were trained on randomized gene assignments to the Hi-Plex data while keeping the test dataset unchanged, thereby disrupting feature–outcome relationships while preserving dataset size and structure. Pearson correlation coefficients were calculated using SciPy [53] between predicted and actual log_2_FC values. Directional prediction accuracy was evaluated through quadrant analysis, categorizing genes based on the signs of predicted and actual log_2_FC values. To assess the statistical significance of the association between predicted and actual expression directions, we first filtered genes with |log_2_FC| ≥ 0.5 based on both the actual gene expression and the predicted gene expression. We then applied Fisher’s exact test to test the association between predicted and actual expression directions for each model and reported the odds ratios and p-values.

To further evaluate each model’s performance in predicting differential expression, after removing genes that are not used for model training as described in the prediction model section, the top 2,000 genes with the largest |log_2_FC| were identified from the RNA-seq data to serve as the true differentially expressed genes. Next, these genes were ranked according to the decreasing |log_2_FC| values calculated based on the predicted gene expression. We evaluated the proportion of the top N predicted differential genes with correctly predicted differential directions according to the actual gene expression (left panel; **Fig. 5D**). Similarly, genes were also ranked according to the decreasing |log_2_FC| values calculated derived from the true gene expression, and we evaluated the proportion of the top N RNA-seq-defined differentially expressed genes with correctly predicted directions of changes (**Fig. 5D**: right panel). Finally, Precision-Recall curves were generated by plotting precision versus recall at different prediction cutoff values (**Supplementary Fig. S6B**).

### TFs and proteins binding site enrichment analysis

To investigate the potential regulatory functions of the combinatorial co-occurrence patterns of the CRF pairs, we performed a binding site enrichment analysis for each genomic region cluster using binding sites of 309 TFs and DNA-binding proteins from ChIP-seq peak datasets of K562 cells provided by ENCODE [20]. Control regions were generated for the genomic regions in all clusters using the refgene_getmatchedcontrol function in CisGenome [54]. Given the regions in a cluster as input regions, for each TF, we calculated the number of TF ChIP-seq peak overlapping with the input regions and control regions, respectively. Enrichment was assessed using Fisher’s exact test, comparing the frequency of overlaps in the input regions versus the control regions for each TF. To account for multiple testing, p-values were adjusted to false discovery rates (FDRs) using the Benjamini-Hochberg (BH) procedure. TFs with total overlap counts below the 10^th^ percentile across all TFs and clusters were excluded to ensure sufficient data quality. Only TFs that met the following two criteria were retained for further analysis and visualization: (1) Statistical significance: FDR < 0.05; (2) Enrichment strength: Odds ratio ≥ 2 in at least one cluster.

### DNA methylation data

Whole-genome bisulfite sequencing (WGBS) DNA methylation data for K562 was downloaded from ENCODE (accession number: ENCSR765JPC) [20].

### Differential Hi-Plex CUT&Tag analysis

To compare Hi-Plex CUT&Tag profiles between sodium butyrate-treated (T) and untreated control (C) K562 cells, we performed differential analyses. To reduce the analysis complexity and noise of individual CRF pairs, the analysis was done on each CRF pair cluster defined in **Fig. 3A**, aggregating all CRF pairs within the cluster. Specifically, for each column cluster in **Fig. 3A**, raw reads were summed up across all CRF pairs within the cluster for each genomic region in each sample (C1, C2, T1, and T2). Genomic regions with total read counts less than 2 were excluded. Genomic regions with non-zero summarized reads in at least one condition and two samples (C1 & C2 or T1 & T2) were retained for differential analysis. The summarized reads of these regions from the four samples were used to compare the two conditions (T vs. C) using the edgeR package [55]. Significance was determined using FDR<0.25 and |log_2_FC| > 0.5. The analysis revealed 456 to 120,829 significantly differential genomic regions between T and C in different CRF clusters except clusters 9 and 16. These results are visualized in **Supplementary Fig. S5D**, where significantly differential genomic regions are displayed as log_2_-transformed read counts, sorted by ascending log_2_FC.

### Visualization of single-cell data

The Hi-Plex CUT&Tag data obtained from K562 single cells using a panel of 16 mAbs and IgG was processed using algorithms similar to those used for the bulk Hi-Plex CUT&Tag data analysis. After data demultiplexing and alignment, cells that contain zero aligned reads were excluded, resulting in 1,278 and 1,318 cells from two replicates, respectively. We then mapped the read counts for each CRF pair in every cell to consecutive 10,000 bp genomic regions across the entire genome. The counts were further log_10_-transformed after adding a pseudocount of 1 to ensure numerical stability. In the bottom panel of **Fig. 6A**, we showed a heatmap for the 100 cells with the highest read counts within the selected region for H3K4me3 homotone and H3K27me3 homotone, while for H3K4me3-H3K27me3, all 88 cells with non-zero reads within the selected region were shown.

### Reproducibility of CRF pair-specific signals between single-cell and bulk data

To evaluate the reproducibility of single-cell data with respect to bulk data for each CRF pair, we aggregated single-cell data into pseudobulk data. For each CRF pair, the pseudobulk data were generated by aggregating the aligned read count files (bam files) of QC-passed cells using the merge function in SAMtools [56]. The pseudobulk read counts were mapped onto the peak regions from the bulk data of the corresponding CRF pairs. We then compared the bulk and pseudobulk data using scatter plots shown in **Fig. 6B**. Pearson correlation coefficients were also computed to quantify the concordance of signals in these peak regions between bulk and pseudobulk data.

### Reproducibility of CRF pair patterns in variable genomic regions

In the bulk data, the CRF pair patterns were visualized as a biclustering heatmap (**Fig. 3A**). To assess the reproducibility of signal patterns from the single-cell data, we focused on 92 CRF pairs that overlapped between single-cell data and the 501 QC-passed CRF pairs in the bulk data. For the single-cell data, we first generated a pseudobulk profile for each CRF pair by aggregating reads across cells. We extracted the pseudobulk signals of the 92 CRF pairs from the same genomic regions in **Fig. 3A**. By keeping the row and column orders the same as in **Fig. 3A**, we then generated the biclustering heatmap for the pseudobulk data and compared it to the heatmap of the bulk data for the same CRF pairs and regions.

### Co-occurrence of two CRF pairs in single cells

We next dissected the co-occurrence patterns of CRF pairs for each 800 bp genomic region from the bulk data down to the single-cell level to examine the co-occurrence of two CRF pairs. We only consider cells whose total reads of all CRF pairs are no less than 1. The “co-occurrence” of two CRF pairs refers to the presence of read counts for both CRF pairs within the same genomic region and cell. We performed permutation tests to determine whether the observed co-occurrence was statistically significant in order to distinguish between randomly observed and biologically driven co-occurrence.

Given two CRF pairs, we first counted their observed co-occurrence, defined as the number of cell-genomic-region combinations with non-zero read counts for both CRF pairs, computed across all cells and the 105,519 genome-wide biclustering analysis regions identified in **Fig. S3A**. Next, we shuffled the cell labels for one of the CRF pairs and recounted the number of co-occurrences with the other CRF pair. This shuffling process was repeated 10,000 times to generate a permutation null distribution of the co-occurrence count. To characterize statistical significance, the observed co-occurrence count was compared to the permutation null distribution, and p-value was calculated as the fraction of permutations where the simulated co-occurrence count was greater than or equal to the observed count. To quantify biological relevance, we also calculated an enrichment score, which was defined as the ratio between the observed co-occurrence count and the mean of simulated co-occurrence counts across the permutations. A pseudocount of 1 was added to both the numerator and denominator to stabilize the results.

Among the 92 CRF pairs overlapping with QC-passed bulk data, we tested only pairs with sufficient observable co-occurrence events, defined as having read counts in more than five cell-genomic-region instances, resulting in 97 combinations of CRF pairs.

The significance and enrichment of co-occurrent CRF pairs are visualized using lower triangle heatmaps. P-values were converted to FDR using the BH method, and-log_10_(FDR) values are displayed. When the permutation p-value was 0 – indicating that all the 10,000 permutations produced a simulated co-occurrence count smaller than the observed count – a pseudocount of 1 was added to both the numerator and denominator to yield a p-value of 1/10,001, avoiding - log_10_(0). For the enrichment score, log_2_(enrichment) values were visualized. The significant co-occurrence events were defined using FDR < 0.05 and log_2_(enrichment) > 1.5, resulting in 76 significant co-occurring CRF pair combinations.

## Notes

### Competing Interest Statement

The authors have declared no competing interest.

## REFERENCES

1. Strahl, B.D. and C.D. Allis, The language of covalent histone modifications. Nature, 2000. 403(6765): p. 41–5.

2. Jenuwein, T. and C.D. Allis, Translating the histone code. Science, 2001. 293(5532): p. 1074–80.

3. Taverna, S.D., et al., How chromatin-binding modules interpret histone modifications: lessons from professional pocket pickers. Nat Struct Mol Biol, 2007. 14(11): p. 1025–1040.

4. Ruthenburg, A.J., et al., Multivalent engagement of chromatin modifications by linked binding modules. Nat Rev Mol Cell Biol, 2007. 8(12): p. 983–94.

5. Klein, D.C. and S.J. Hainer, Genomic methods in profiling DNA accessibility and factor localization. Chromosome Res, 2020. 28(1): p. 69–85.

6. Park, P.J., ChIP-seq: advantages and challenges of a maturing technology. Nat Rev Genet, 2009. 10(10): p. 669–80.

7. Zentner, G.E. and S. Henikoff, High-resolution digital profiling of the epigenome. Nat Rev Genet, 2014. 15(12): p. 814–27.

8. Skene, P.J. and S. Henikoff, An efficient targeted nuclease strategy for high-resolution mapping of DNA binding sites. Elife, 2017. 6.

9. Kaya-Okur, H.S., et al., *CUT&*Tag for efficient epigenomic profiling of small samples and single cells. Nat Commun, 2019. 10(1): p. 1930.

10. Meers, M.P., et al., Multifactorial profiling of epigenetic landscapes at single-cell resolution using MulTI-Tag. Nat Biotechnol, 2023. 41(5): p. 708–716.

11. Xiong, H., et al., Single-cell joint profiling of multiple epigenetic proteins and gene transcription. Sci Adv, 2024. 10(1): p. eadi3664.

12. Lochs, S.J.A., et al., Combinatorial single-cell profiling of major chromatin types with MAbID. Nat Methods, 2024. 21(1): p. 72–82.

13. Perez, A.A., et al., ChIP-DIP maps binding of hundreds of proteins to DNA simultaneously and identifies diverse gene regulatory elements. Nat Genet, 2024. 56(12): p. 2827–2841.

14. Gopalan, S. and T.G. Fazzio, Multi-CUT&Tag to simultaneously profile multiple chromatin factors. STAR Protoc, 2022. 3(1): p. 101100.

15. Gopalan, S., et al., Simultaneous profiling of multiple chromatin proteins in the same cells. Mol Cell, 2021. 81(22): p. 4736–4746 e5.

16. Stuart, T., et al., Nanobody-tethered transposition enables multifactorial chromatin profiling at single-cell resolution. Nat Biotechnol, 2023. 41(6): p. 806–812.

17. Barcenas-Walls, J.R., et al., Nano-CUT&Tag for multimodal chromatin profiling at single-cell resolution. Nat Protoc, 2024. 19(3): p. 791–830.

18. Bartosovic, M. and G. Castelo-Branco, Multimodal chromatin profiling using nanobody-based single-cell CUT&Tag. Nat Biotechnol, 2023. 41(6): p. 794–805.

19. Green, N.M., Avidin. Adv Protein Chem, 1975. 29: p. 85–133.

20. Consortium, E.P., An integrated encyclopedia of DNA elements in the human genome. Nature, 2012. 489(7414): p. 57–74.

21. Consortium, E.P., et al., Expanded encyclopaedias of DNA elements in the human and mouse genomes. Nature, 2020. 583(7818): p. 699–710.

22. Smit, A., Hubley, R & Green, P. RepeatMasker Open-4.0. 2013-2015; Available from: <http://www.repeatmasker.org>.

23. Kulpa, D.A. and J.V. Moran, Ribonucleoprotein particle formation is necessary but not sufficient for LINE-1 retrotransposition. Hum Mol Genet, 2005. 14(21): p. 3237–48.

24. Iwamoto, S., et al., Cloning and characterization of erythroid-specific DNase I-hypersensitive site in human rhesus-associated glycoprotein gene. J Biol Chem, 2000. 275(35): p. 27324–31.

25. Jones, P.A., Functions of DNA methylation: islands, start sites, gene bodies and beyond. Nat Rev Genet, 2012. 13(7): p. 484–92.

26. Ernst, J. and M. Kellis, Chromatin-state discovery and genome annotation with ChromHMM. Nat Protoc, 2017. 12(12): p. 2478–2492.

27. Patton, J.G., et al., Cloning and characterization of PSF, a novel pre-mRNA splicing factor. Genes Dev, 1993. 7(3): p. 393–406.

28. Mathur, M., P.W. Tucker, and H.H. Samuels, PSF is a novel corepressor that mediates its effect through Sin3A and the DNA binding domain of nuclear hormone receptors. Mol Cell Biol, 2001. 21(7): p. 2298–311.

29. Bertolotti, A., et al., EWS, but not EWS-FLI-1, is associated with both TFIID and RNA polymerase II: interactions between two members of the TET family, EWS and hTAFII68, and subunits of TFIID and RNA polymerase II complexes. Mol Cell Biol, 1998. 18(3): p. 1489–97.

30. Bertolotti, A., et al., hTAF(II)68, a novel RNA/ssDNA-binding protein with homology to the pro-oncoproteins TLS/FUS and EWS is associated with both TFIID and RNA polymerase II. Embo j, 1996. 15(18): p. 5022–31.

31. Chen, C.D., R. Kobayashi, and D.M. Helfman, Binding of hnRNP H to an exonic splicing silencer is involved in the regulation of alternative splicing of the rat beta-tropomyosin gene. Genes Dev, 1999. 13(5): p. 593–606.

32. Gabler, S., et al., E1B 55-kilodalton-associated protein: a cellular protein with RNA-binding activity implicated in nucleocytoplasmic transport of adenovirus and cellular mRNAs. J Virol, 1998. 72(10): p. 7960–71.

33. Ferreira, R., et al., GATA1 function, a paradigm for transcription factors in hematopoiesis. Mol Cell Biol, 2005. 25(4): p. 1215–27.

34. Vicente, C., et al., The role of the GATA2 transcription factor in normal and malignant hematopoiesis. Crit Rev Oncol Hematol, 2012. 82(1): p. 1–17.

35. Nosaka, T., et al., STAT5 as a molecular regulator of proliferation, differentiation and apoptosis in hematopoietic cells. Embo j, 1999. 18(17): p. 4754–65.

36. Luo, H., et al., The functions of SET domain bifurcated histone lysine methyltransferase 1 (SETDB1) in biological process and disease. Epigenetics Chromatin, 2023. 16(1): p. 47.

37. Schoelz, J.M. and N.C. Riddle, Functions of HP1 proteins in transcriptional regulation. Epigenetics Chromatin, 2022. 15(1): p. 14.

38. Noreen, S., et al., Unravelling the impact of the chromobox proteins in human cancers. Cell Death Dis, 2025. 16(1): p. 238.

39. Forsburg, S.L., Eukaryotic MCM proteins: beyond replication initiation. Microbiol Mol Biol Rev, 2004. 68(1): p. 109–31.

40. Breiman, L., Random Forests. Machine Learning, 2001. 45(1): p. 5–32.

41. Davie, J.R., Inhibition of histone deacetylase activity by butyrate. J Nutr, 2003. 133(7 Suppl): p. 2485s–2493s.

42. Song, G., et al., An all-to-all approach to the identification of sequence-specific readers for epigenetic DNA modifications on cytosine. Nat Commun, 2021. 12(1): p. 795.

43. Thermo Fisher Scientific, Multiple Primer Analyzer.; Available from: https://www.thermofisher.com/us/en/home/brands/thermo-scientific/molecular-biology/molecular-biology-learning-center/molecular-biology-resource-library/thermo-scientific-web-tools/multiple-primer-analyzer.html.

44. Martin, M., Cutadapt removes adapter sequences from high-throughput sequencing reads. EMBnet.journal, 2011. v. 17, n. 1, p. pp. 10–12, may 2011.

45. Royer, C.A., et al., Ligand binding and protein dynamics: a fluorescence depolarization study of aspartate transcarbamylase from Escherichia coli. Biochemistry, 1987. 26(20): p. 6472–8.

46. Langmead, B. and S.L. Salzberg, Fast gapped-read alignment with Bowtie 2. Nat Methods, 2012. 9(4): p. 357–9.

47. Gu, Z., R. Eils, and M. Schlesner, Complex heatmaps reveal patterns and correlations in multidimensional genomic data. Bioinformatics, 2016. 32(18): p. 2847–9.

48. Meers, M.P., D. Tenenbaum, and S. Henikoff, Peak calling by Sparse Enrichment Analysis for CUT&RUN chromatin profiling. Epigenetics Chromatin, 2019. 12(1): p. 42.

49. Yu, G., L.G. Wang, and Q.Y. He, ChIPseeker: an R/Bioconductor package for ChIP peak annotation, comparison and visualization. Bioinformatics, 2015. 31(14): p. 2382–3.

50. Lawrence, M., et al., Software for computing and annotating genomic ranges. PLoS Comput Biol, 2013. 9(8): p. e1003118.

51. Fabian Pedregosa, G.V., Alexandre Gramfort, Vincent Michel, Bertrand Thirion, Olivier Grisel, Mathieu Blondel, Peter Prettenhofer, Ron Weiss, Vincent Dubourg, Jake Vanderplas, Alexandre Passos, David Cournapeau, Matthieu Brucher, Matthieu Perrot, Édouard Duchesnay, Scikit-learn: Machine Learning in Python. Journal of Machine Learning Research, 2011. 12: p. 2825–2830.

52. Satopaa, V., et al., Finding a Kneedle in a Haystack: Detecting Knee Points in System Behavior. 2011. 166–171.

53. Virtanen, P., et al., SciPy 1.0: fundamental algorithms for scientific computing in Python. Nat Methods, 2020. 17(3): p. 261–272.

54. Ji, H., et al., Using CisGenome to analyze ChIP-chip and ChIP-seq data. Curr Protoc Bioinformatics, 2011. **Chapter 2**: p. Unit2 13.

55. Robinson, M.D., D.J. McCarthy, and G.K. Smyth, edgeR: a Bioconductor package for differential expression analysis of digital gene expression data. Bioinformatics, 2010. 26(1): p. 139–40.

56. Danecek, P., et al., Twelve years of SAMtools and BCFtools. Gigascience, 2021. 10(2).

