## Supplementary material for "Global Mapping of Combinatorial Chromatin Regulatory Events Using Hi-Plex CUT&Tag": Hi-Plex CUT&Tag Supplementary materials

### Supplementary notes

#### Differential Hi-Plex CUT&Tag analysis between treated and untreated K562 cells

Consistent with the known effect of sodium butyrate, the DNA fragment length distribution shifted towards shorter fragments following treatment (**Supplementary Fig. S5B**). Of the 501 CRF pairs analyzed, nearly all (495 pairs) showed an increased proportion of sub-nucleosomal fragments after treatment, with 247 pairs exhibiting statistically significant positive changes (Fisher's exact test, FDR < 0.05). Concurrently, 426 of the 501 CRF pairs demonstrated a significant decrease in the proportion of di- and multi-nucleosomal fragments. In contrast, we observed minimal changes in the proportion of mono-nucleosomal fragments, with only 3 CRF pairs showing significant differences. These results reflect chromatin relaxation induced by sodium butyrate, leading to a shift toward shorter fragment lengths.

We also conducted a differential analysis to examine changes in CRF co-localization signals after treatment. To simplify presentation and increase statistical power, we aggregated data across all CRF pairs within each cluster (defined in **Fig. 3A**) and performed the analysis for each CRF pair cluster. Fourteen of the 16 CRF pair clusters showed significant changes (**Supplementary Fig. S5C**). Most clusters displayed a higher number of up-regulated regions (i.e., regions with higher Hi-Plex signals in treated samples) than down-regulated regions (i.e., regions with higher Hi-Plex signals in control samples) following treatment. For example, Cluster 1, which contains CRF pairs involving H3K4me3, showed increased signals in 19,718 regions and decreased signals in 7,779 regions (**Supplementary Fig. S5D**). Some clusters exhibited a more balanced distribution of up- and down-regulated regions. For instance, Cluster 8, which is enriched in H3K27ac-associated pairs, including bivalent combinations like H3K27ac-H3K9me3, showed 5,570 up-regulated and 5,705 down-regulated regions (**Supplementary Fig. S5D**).

Further exploration of the relationship between $log_2FC$ of Hi-plex CUT&Tag signal of each CRF pair cluster and differential gene expression was conducted by mapping the differential Hi-Plex regions to the promoter regions of genes (5000 bp upstream of TSS to TES). For genes whose promoter region overlapped with multiple significantly differential Hi-Plex regions, we summarized the Hi-Plex signal by taking the average $log_2FC$ across these regions and assigned the minimum FDR among them as the gene-level Hi-Plex FDR. Gene expression $log_2FC$ was computed based on the average TPM across conditions with a pseudocount of 1, and the corresponding FDR was assessed using the DESeq2 R package [1]. For each column cluster, a scatter plot (**Supplementary Fig S5E**) was generated with $log_2FC$ of gene expression on the Y-axis and $log_2FC$ of Hi-plex CUT&Tag signal on the X-axis. Significant observations, defined by $|log_2FC| > 0.5$ and $FDR < 0.25$ for both axes, were highlighted and a trend line was fitted using the geom_smooth function in ggplot2 [2]. Fisher's exact test (base R function) was performed on the number of significant observations in each quadrant to calculate Odds Ratios (OR) and corresponding p-values for each column cluster.

### Supplementary figures

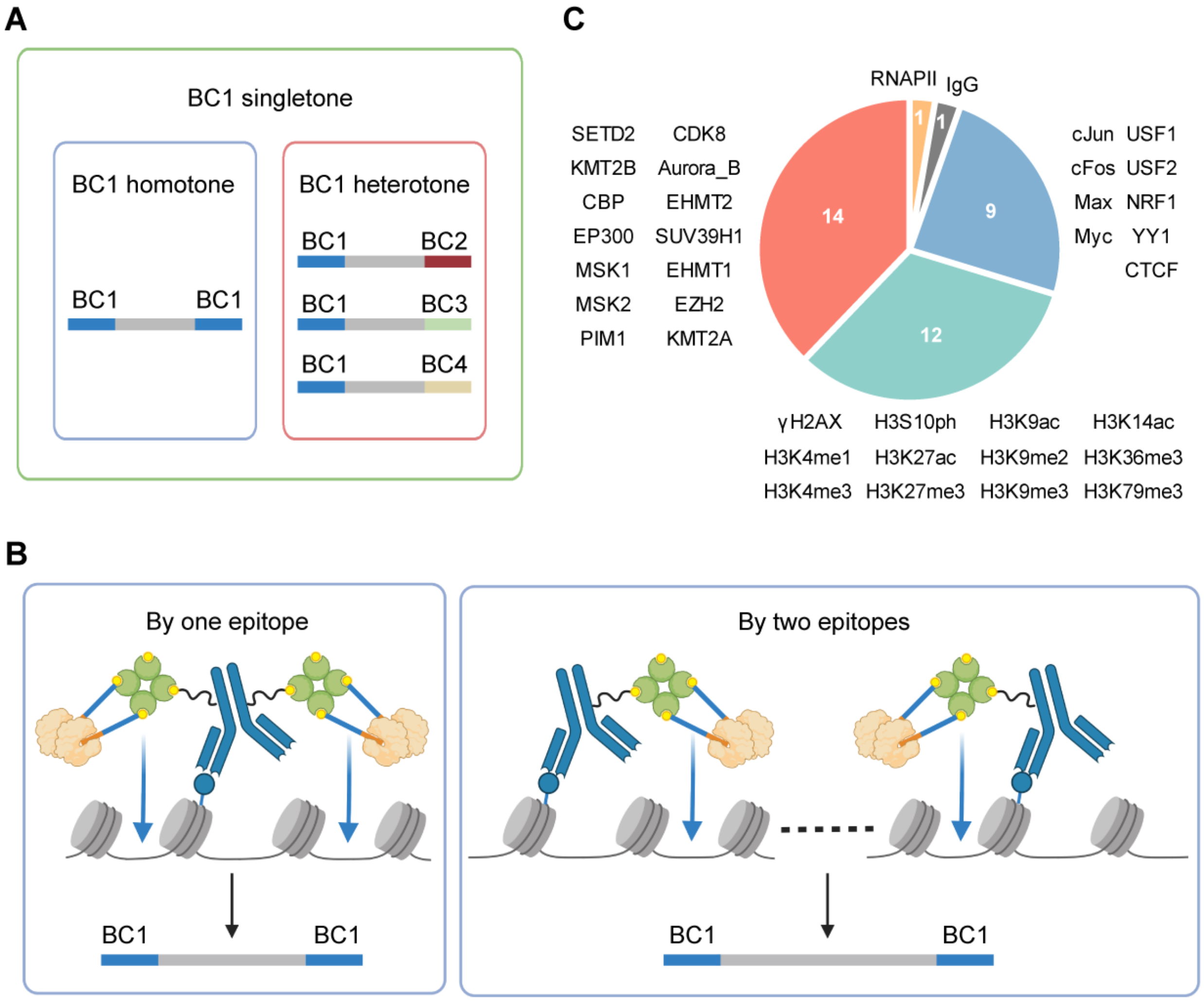


**Figure S1. Summary of the 37 barcoded antibodies used in Hi-Plex CUT&Tag.**

**A.** Cartoon illustrations defining singletone, homotone, and hererotone. **B.** Two scenarios which will produce homotone fragments. **C.** We barcoded a panel of 37 Abs, targeting 12 common histone marks (light green), 14 histone modification enzymes (salmon), nine human TFs (light blue), PolII (pSer2) (orange), and Rabbit IgG negative control (gray), respectively.

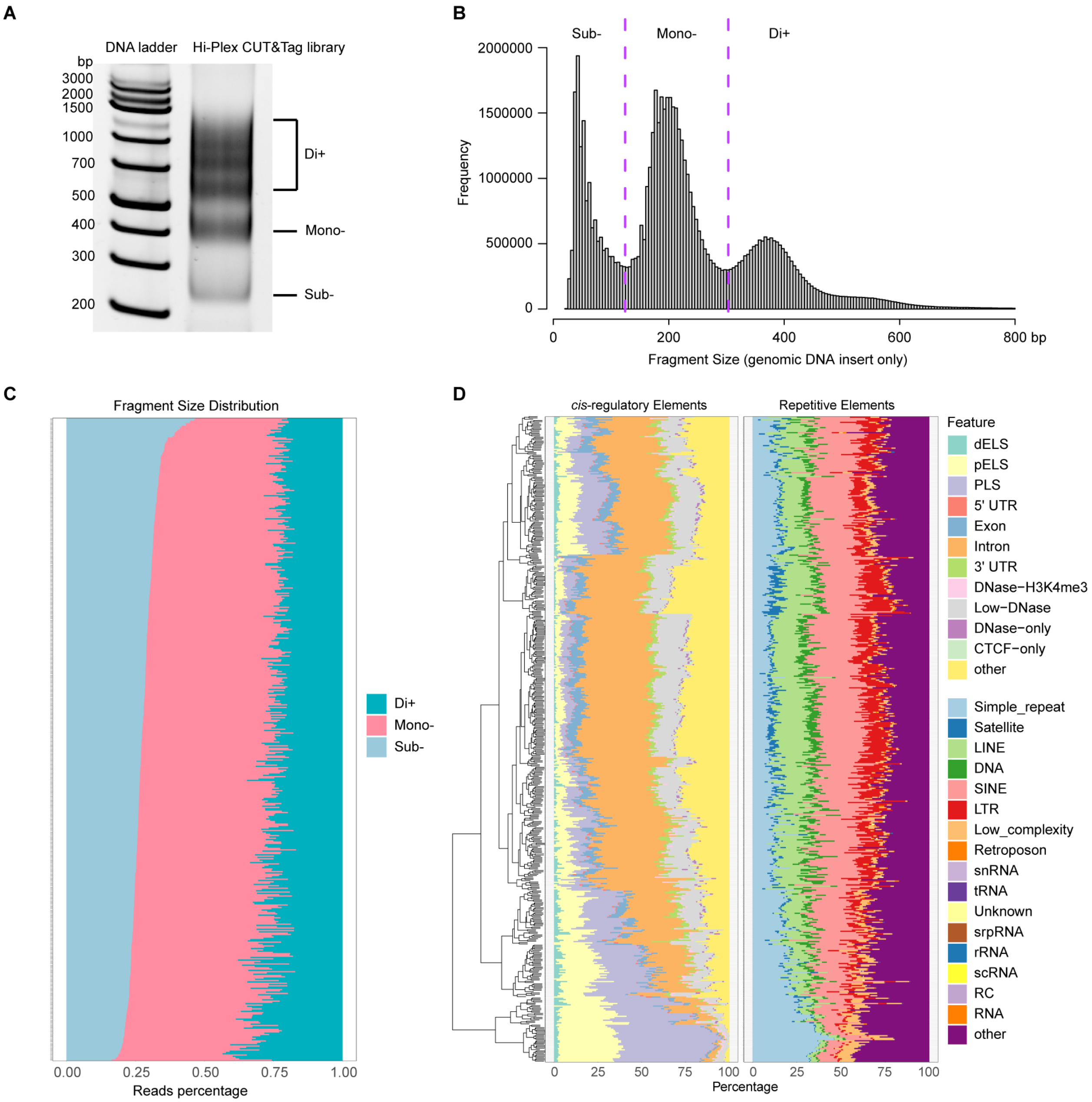


**Figure S2. Size distribution of Hi-Plex CUT & Tag fragments**

**Figure S2. Size distribution of Hi-Plex CUT & Tag fragments**

**A.** Gel picture showing the laddering pattern of Hi-Plex CUT&Tag library. Different size of the whole fragments including adaptors and barcodes is labeled as sub -, mono -, di+ (relating to the number of nucleosomes occupying the endogenous fragment). **B.** Analysis of size distribution of all the fragments without adaptors and barcodes from Hi-Plex CUT & Tag library. Names of different sizes are labeled on the top of each peak. **C.** Fragment length stacked bar plot for all CRF pairs sorted in increasing sub-nucleosome distance proportion. **D.** cis-regulatory elements and repetitive elements annotation of all CRF pairs.

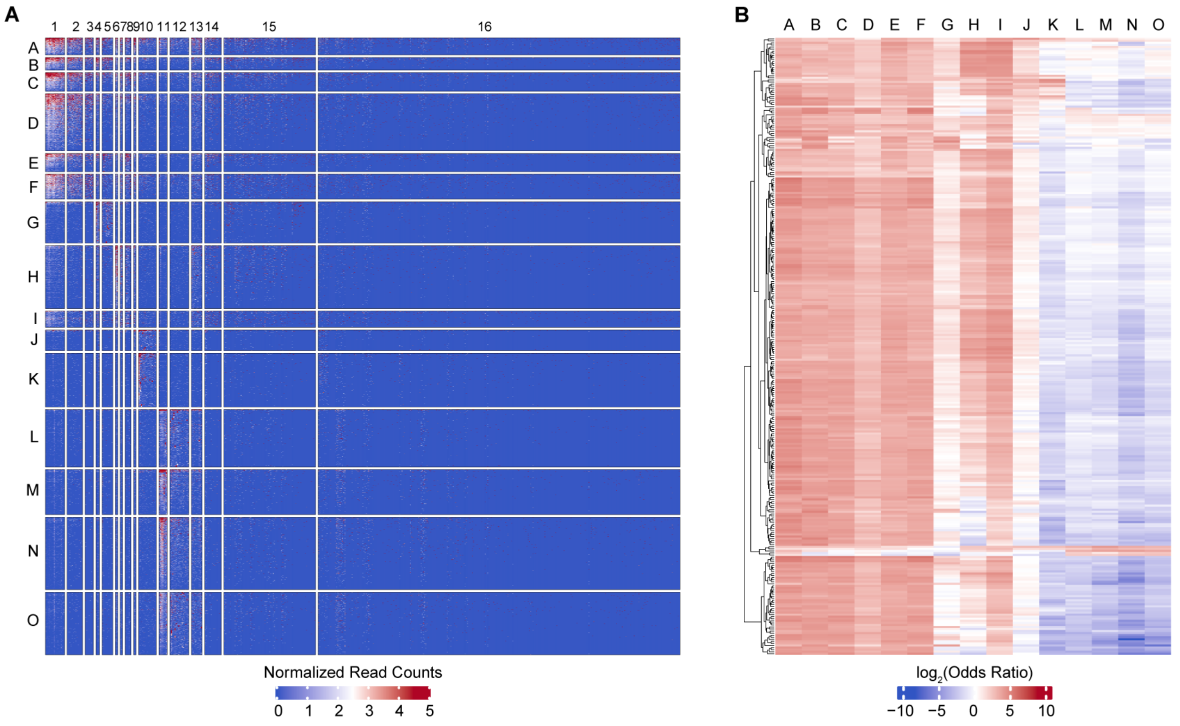


**Figure S3. Analysis of the combinatorial landscape of co-localized CRF pairs.**

**A.** Heatmap of the full set of genomic regions. **B.** Heatmap showing the odds ratio from the ChIP-Seq peak enrichment analysis.

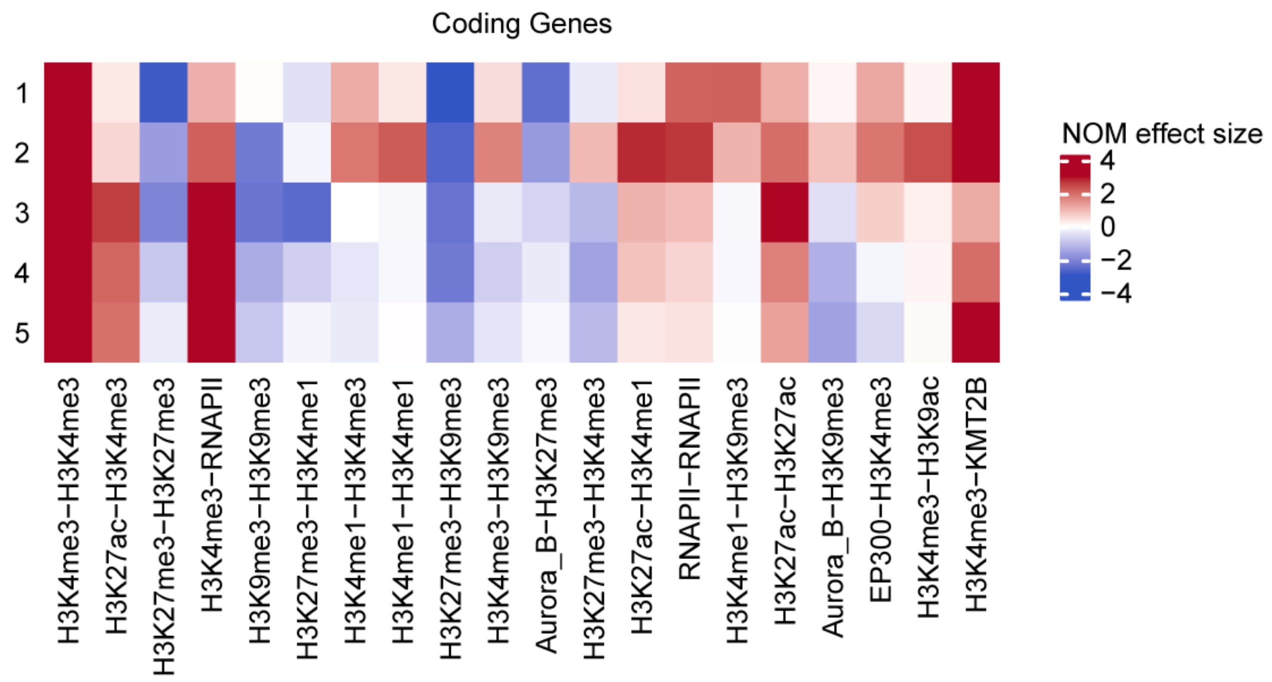


**Figure S4. Effect size representing the signed area under the partial dependence curve for the top 20 CRF pairs from the Random Forest model.**

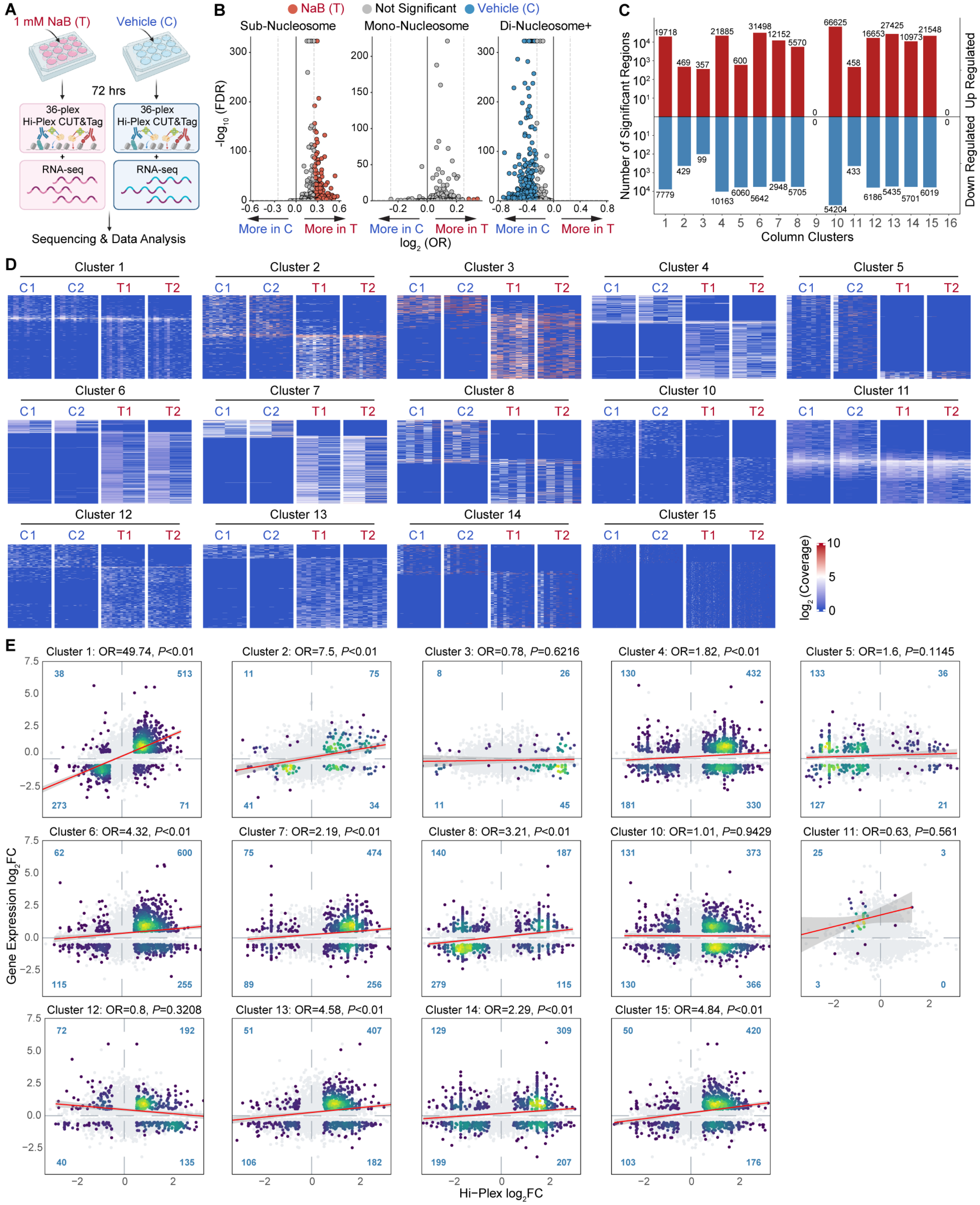


**Figure S5. Differential Hi-Plex CUT&Tag signals between the treated and untreated samples in different CRF pair clusters and their various relationship with gene expression.**

**Figure S5. Differential Hi-Plex CUT&Tag signals between the treated and untreated samples in different CRF pair clusters and their various relationship with gene expression.**

**A.** Global chromatin remodeling induced with sodium butyrate treatment in K562 cells. Hi-Plex CUT&Tag and RNA-seq were performed in parallel on both treated (T) and untreated (C) samples. **B.** Volcano plots showing treatment-induced changes in proportions of sub-nucleosome, mono-nucleosome, and di-nucleosome+ fragments for each CRF pair. Blue indicates significantly higher proportions before treatment (FDR < 0.05, $\log_2$FC < -0.25), red indicates significantly higher after treatment (FDR < 0.05, $\log_2$FC > 0.25), and grey indicates no significant change. **C.** Bar plot showing the number of genomic regions which showed significant differential Hi-Plex signals (FDR < 0.25) between treated and untreated samples for each column cluster defined in **Fig. 3A.** Red indicates higher Hi-Plex CUT&Tag signals after treatment, and blue indicates higher signals before treatment. **D.** Heatmaps comparing the $\log_2$-transformed Hi-Plex CUT&Tag signals at the significantly differential genomic regions for CRF pairs in each of the 15 column clusters, defined in **Fig. 3A**, between the treated (T1 and T2) and untreated samples (C1 and C2). **E.** Scatterplots showing the various relationships between the change of gene expression and the change of Hi-plex CUT&Tag signals in each CRF pair cluster.

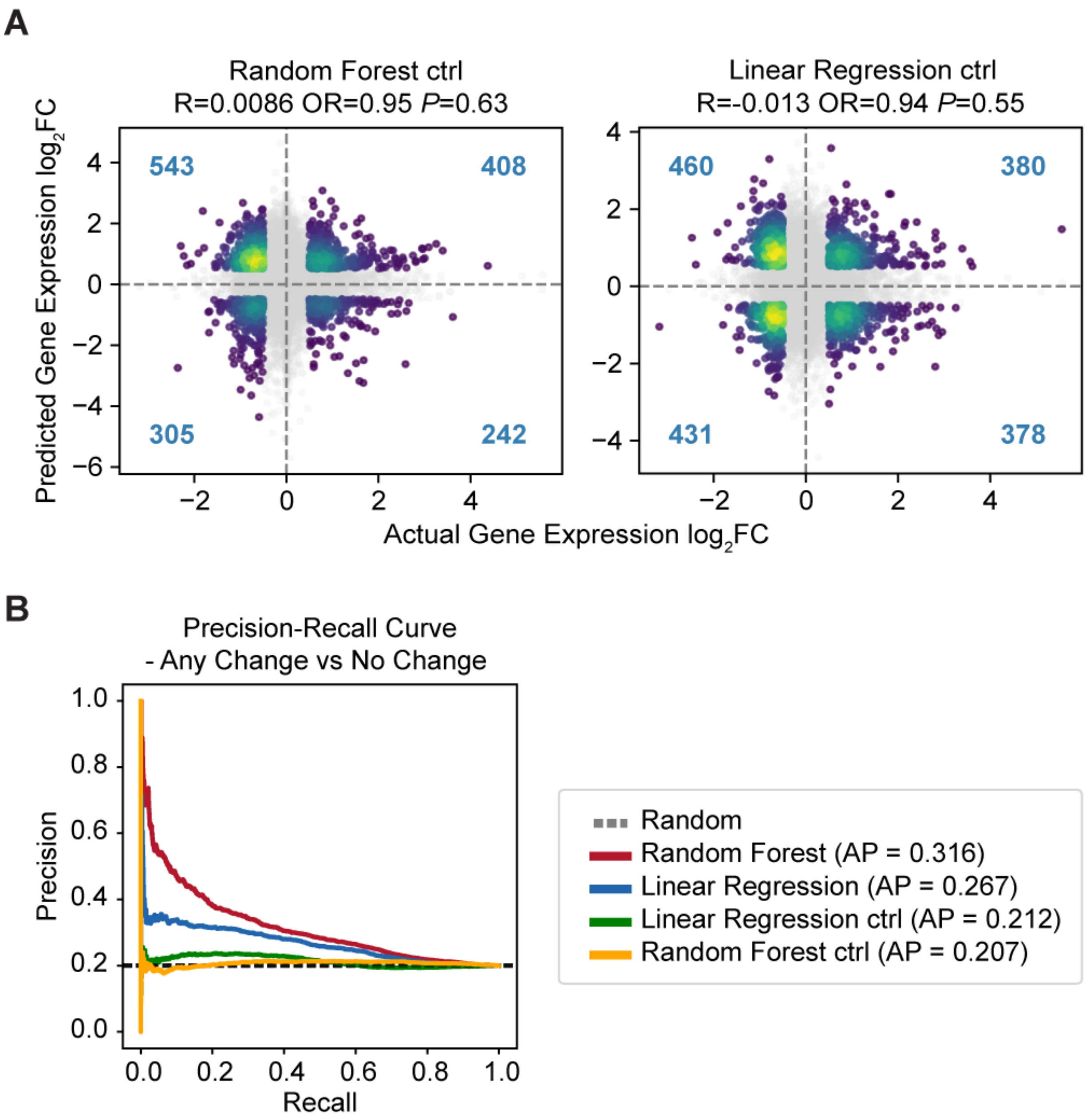


**Figure S6. Dynamic changes in Hi-Plex CUT&Tag predict gene expression changes.**

**A.** Scatterplots comparing the true and predicted $\log_2$ fold change of gene expression between treated and untreated samples for the control models based on Random Forest and Linear Regression, respectively. **B**. Precision-recall curves comparing the performance of Random Forest, Linear Regression, and the two control models for predicting the gene expression changes based on Hi-Plex signals.

### Supplementary tables

#### Table S1. Annotation Category and Description.

The Category column represents the labels shown in the annotation stacked barplot. The description column represents the functional definition of the corresponding category.

| category | description |
|---|---|
| **ENCODE cCREs** | |
| dELS | distal enhancer-like signature |
| pELS | proximal enhancer-like signature |
| PLS | promoter-like signatures |
| 5' UTR | 5 end untranslated regions |
| Exon | exon |
| Intron | intron |
| 3' UTR | 3 end untranslated regions |
| Dnase-H3K4me3 | cCREs have high H3K4me3 max-Z scores but low H3K27ac max-Z scores and do not fall within 200 bp of a TSS. |
| Dnase only | cCREs with high DNase Z-scores but low H3K4me3, H3K27ac, and CTCF Z-scores |
| CTCF only | CTCF-only cCREs have high DNase and CTCF max-Z scores and low H3K4me3 and H3K27ac max-Z scores. |
| **Repetitive elements** | |
| Simple Repeat | micro-satellites |
| Satellite | Satellite repeats |
| LINE | Long interspersed nuclear elements |
| DNA repeats | DNA repeat elements |
| SINE | Short interspersed nuclear elements |
| LTR | Long terminal repeat elements |
| Low Complexity | Low complexity repeats |
| Retroposon | Retroposon |
| snRNA | snRNA repeats |
| tRNA | tRNA repeats |

| | |
|---|---|
| Unknown | Unknown repeats |
| srpRNA | srpRNA repeats |
| rRNA | rRNA repeats |
| scRNA | scRNA repeats |
| RC | Rolling Circle |
| RNA | RNA repeats |
| **ChromHMM** | |
| GapArtf | Assembly gaps and artifacts |
| Acet | acetylations |
| EnhWk | weak enhancers |
| Quies | quescient |
| TxWk | weak transcription |
| TxEX | exon |
| EnhA | enhancers |
| ReprRC | polycomb repressed |
| HET | HET |
| Dnase | DNase |
| PromF | promoters |
| Tx | transcription |
| TSS | TSS |
| znf | znf |
| TxEnh | transcribed and enhancer |
| BivProm | weak promoters |

**Table S2. CRF pairs in each column cluster**

| CRF_pairs | Cluster | CRF_pairs | Cluster | CRF_pairs | Cluster | CRF_pairs | Cluster |
|---|---|---|---|---|---|---|---|
| H3K4me3-RNAPII | 1 | MSK1-RNAPII | 15 | EP300-H3K79me3 | 16 | H3S10ph-MSK1 | 16 |
| H3K27ac-H3K4me3 | 1 | RNAPII-cJun | 15 | EP300-γH2AX | 16 | EZH2-H3K9me2 | 16 |
| H3K4me3-H3K4me3 | 1 | KMT2A-RNAPII | 15 | EP300-MSK2 | 16 | H3K36me3-H3S10ph | 16 |

| | | | | | | | |
|---|---|---|---|---|---|---|---|
| H3K4me1-H3K4me3 | 1 | H3K4me1-H3S10ph | 15 | CDK8-H3K36me3 | 16 | H3K36me3-PIM1 | 16 |
| H3K4me3-H3K9ac | 1 | H3K4me1-H3K79me3 | 15 | CDK8-H3S10ph | 16 | H3K9me2-USF2 | 16 |
| EP300-H3K4me3 | 1 | H3K4me1-PIM1 | 15 | CBP-EZH2 | 16 | EHMT1-H3K9me2 | 16 |
| H3K4me3-H3K9me3 | 1 | γH2AX-H3K4me1 | 15 | CBP-PIM1 | 16 | H3S10ph-KMT2A | 16 |
| H3K27me3-H3K4me3 | 1 | H3K4me1-KMT2A | 15 | PIM1-SUV39H1 | 16 | CBP-USF1 | 16 |
| H3K4me3-SUV39H1 | 1 | KMT2B-KMT2B | 15 | EP300-Myc | 16 | KMT2B-Max | 16 |
| H3K4me3-KMT2B | 1 | SETD2-SETD2 | 15 | CBP-KMT2A | 16 | Aurora_B-YY1 | 16 |
| CBP-H3K4me3 | 1 | H3K9ac-KMT2B | 15 | CTCF-SETD2 | 16 | SETD2-YY1 | 16 |
| CDK8-H3K4me3 | 1 | CTCF-H3K9ac | 15 | EZH2-KMT2B | 16 | H3K9me2-cFos | 16 |
| CTCF-H3K4me3 | 1 | CBP-H3K9ac | 15 | γH2AX-SUV39H1 | 16 | H3K14ac-H3S10ph | 16 |
| Aurora_B-H3K4me3 | 1 | CDK8-H3K9ac | 15 | H3K9me3-cFos | 16 | γH2AX-H3K36me3 | 16 |
| H3K4me3-H3K9me2 | 1 | H3K9ac-SETD2 | 15 | H3K9me3-YY1 | 16 | EHMT2-cJun | 16 |
| H3K4me3-SETD2 | 1 | EP300-SUV39H1 | 15 | H3K9me3-USF1 | 16 | γH2AX-MSK2 | 16 |
| H3K4me3-KMT2A | 2 | EZH2-H3K4me1 | 15 | KMT2B-MSK2 | 16 | H3S10ph-Myc | 16 |
| H3K4me3-MSK1 | 2 | H3K4me1-MSK1 | 15 | γH2AX-KMT2B | 16 | EHMT1-EHMT1 | 16 |
| H3K4me3-cJun | 2 | H3K4me1-cFos | 15 | CBP-USF2 | 16 | EHMT2-H3K9me2 | 16 |

| | | | | | | | |
|---|---|---|---|---|---|---|---|
| H3K4me3-H3S10ph | 2 | H3K14ac-H3K4me1 | 15 | H3S10ph-SETD2 | 16 | USF1-USF1 | 16 |
| H3K4me3-PIM1 | 2 | CDK8-EP300 | 15 | MSK1-SUV39H1 | 16 | H3S10ph-MSK2 | 16 |
| γH2AX-H3K4me3 | 2 | CBP-EP300 | 15 | KMT2B-MSK1 | 16 | Aurora_B-Max | 16 |
| H3K4me3-MSK2 | 2 | H3K4me1-NRF1 | 15 | Aurora_B-CTCF | 16 | H3K79me3-H3S10ph | 16 |
| H3K36me3-H3K4me3 | 2 | Aurora_B-H3K27ac | 15 | Aurora_B-EZH2 | 16 | MSK2-Myc | 16 |
| H3K4me3-H3K79me3 | 2 | γH2AX-H3K27ac | 15 | Aurora_B-PIM1 | 16 | CBP-Max | 16 |
| EZH2-H3K4me3 | 2 | EP300-KMT2B | 15 | H3K36me3-SETD2 | 16 | γH2AX-MSK1 | 16 |
| H3K14ac-H3K4me3 | 2 | EP300-SETD2 | 15 | SUV39H1-USF2 | 16 | γH2AX-Myc | 16 |
| H3K4me3-Myc | 2 | CTCF-EP300 | 15 | USF2-USF2 | 16 | H3K9me2-YY1 | 16 |
| H3K4me3-cFos | 2 | CDK8-CDK8 | 15 | CBP-H3K36me3 | 16 | H3K9me2-USF1 | 16 |
| H3K4me3-USF2 | 3 | CDK8-KMT2B | 15 | CTCF-KMT2B | 16 | H3K9me2-Max | 16 |
| H3K4me3-YY1 | 3 | CDK8-SUV39H1 | 15 | CTCF-SUV39H1 | 16 | H3K14ac-H3K36me3 | 16 |
| H3K4me3-USF1 | 3 | Aurora_B-H3K9ac | 15 | H3K27me3-USF1 | 16 | γH2AX-H3K79me3 | 16 |
| H3K4me3-NRF1 | 3 | H3K4me1-Myc | 15 | Aurora_B-SETD2 | 16 | EZH2-γH2AX | 16 |
| EHMT1-H3K4me3 | 3 | CBP-SUV39H1 | 15 | H3K27me3-YY1 | 16 | YY1-YY1 | 16 |
| EHMT2-H3K4me3 | 3 | EP300-H3K9me2 | 15 | H3K27me3-Max | 16 | H3K36me3-MSK1 | 16 |

| | | | | | | | |
|---|---|---|---|---|---|---|---|
| H3K4me3-Max | 3 | EHMT1-H3K4me1 | 15 | Aurora_B-cJun | 16 | H3K79me3-MSK2 | 16 |
| RNAPII-RNAPII | 4 | EHMT2-H3K4me1 | 15 | H3K36me3-KMT2B | 16 | EZH2-MSK2 | 16 |
| H3K4me1-RNAPII | 4 | H3K4me1-YY1 | 15 | H3K36me3-SUV39H1 | 16 | Max-SUV39H1 | 16 |
| H3K9me3-RNAPII | 4 | CBP-CDK8 | 15 | H3K14ac-SUV39H1 | 16 | γH2AX-KMT2A | 16 |
| H3K27me3-RNAPII | 5 | EHMT1-RNAPII | 15 | H3K14ac-KMT2B | 16 | SUV39H1-USF1 | 16 |
| H3K27ac-RNAPII | 5 | RNAPII-YY1 | 15 | CBP-γH2AX | 16 | CTCF-MSK1 | 16 |
| H3K9ac-RNAPII | 5 | H3K9ac-PIM1 | 15 | H3K9ac-YY1 | 16 | CTCF-cFos | 16 |
| RNAPII-SUV39H1 | 5 | H3K4me1-MSK2 | 15 | CDK8-MSK1 | 16 | cFos-cJun | 16 |
| CBP-RNAPII | 5 | H3K36me3-H3K4me1 | 15 | CDK8-cFos | 16 | CDK8-EHMT1 | 16 |
| KMT2B-RNAPII | 5 | H3K4me1-Max | 15 | CTCF-cJun | 16 | H3K79me3-H3K9me2 | 16 |
| EP300-RNAPII | 5 | H3K4me1-USF2 | 15 | H3S10ph-KMT2B | 16 | H3K9me2-MSK1 | 16 |
| CDK8-RNAPII | 5 | H3K9ac-H3K9me2 | 15 | KMT2B-cFos | 16 | H3S10ph-H3S10ph | 16 |
| RNAPII-SETD2 | 5 | γH2AX-H3K9ac | 15 | Myc-SUV39H1 | 16 | EZH2-cJun | 16 |
| H3K4me1-H3K4me1 | 6 | H3K36me3-H3K9ac | 15 | CBP-cFos | 16 | Aurora_B-USF1 | 16 |
| CDK8-H3K4me1 | 6 | H3K27ac-H3K79me3 | 15 | SUV39H1-cFos | 16 | H3K14ac-H3K9me2 | 16 |
| H3K4me1-H3K9ac | 7 | H3K27ac-KMT2A | 15 | CBP-H3K79me3 | 16 | H3K36me3-H3K9me2 | 16 |

| | | | | | | | |
|---|---|---|---|---|---|---|---|
| EP300-H3K4me1 | 7 | H3K9me2-RNAPII | 15 | H3K79me3-SUV39H1 | 16 | H3K36me3-MSK2 | 16 |
| H3K27ac-H3K9me3 | 8 | EZH2-RNAPII | 15 | CBP-H3K14ac | 16 | Aurora_B-NRF1 | 16 |
| H3K27ac-H3K4me1 | 8 | H3K36me3-RNAPII | 15 | Aurora_B-H3K9me2 | 16 | H3K9me2-NRF1 | 16 |
| H3K27ac-H3K27ac | 8 | MSK2-RNAPII | 15 | H3K9me2-KMT2B | 16 | NRF1-SETD2 | 16 |
| H3K27ac-H3K27me3 | 8 | H3K14ac-RNAPII | 15 | H3K9me3-Max | 16 | H3K79me3-cJun | 16 |
| H3K27ac-H3K9ac | 8 | Myc-RNAPII | 15 | Aurora_B-H3S10ph | 16 | CTCF-H3K14ac | 16 |
| H3K4me1-H3K9me3 | 9 | H3K79me3-RNAPII | 15 | Aurora_B-H3K79me3 | 16 | γH2AX-PIM1 | 16 |
| H3K9ac-H3K9me3 | 9 | RNAPII-cFos | 15 | SUV39H1-SUV39H1 | 16 | H3K14ac-cJun | 16 |
| H3K9me3-H3K9me3 | 10 | γH2AX-RNAPII | 15 | CBP-EHMT2 | 16 | EZH2-H3K36me3 | 16 |
| Aurora_B-H3K9me3 | 10 | RNAPII-USF2 | 15 | Aurora_B-γH2AX | 16 | EZH2-H3S10ph | 16 |
| H3K9me2-H3K9me3 | 10 | EHMT2-RNAPII | 15 | Aurora_B-MSK1 | 16 | CDK8-NRF1 | 16 |
| H3K9me3-KMT2B | 10 | H3K9ac-cJun | 15 | CTCF-CTCF | 16 | Aurora_B-EHMT1 | 16 |
| H3K9me3-SUV39H1 | 10 | KMT2A-KMT2A | 15 | EHMT2-KMT2B | 16 | MSK2-PIM1 | 16 |
| CBP-H3K9me3 | 10 | H3K9ac-KMT2A | 15 | H3K36me3-H3K36me3 | 16 | EHMT1-SETD2 | 16 |
| H3K9me3-SETD2 | 10 | H3K27ac-H3S10ph | 15 | KMT2B-Myc | 16 | cFos-cFos | 16 |
| CTCF-H3K9me3 | 10 | H3K27ac-MSK1 | 15 | γH2AX-SETD2 | 16 | SUV39H1-YY1 | 16 |

| | | | | | | | |
|---|---|---|---|---|---|---|---|
| H3K9me3-cJun | 10 | H3K27ac-MSK2 | 15 | H3K9me2-cJun | 16 | MSK1-PIM1 | 16 |
| γH2AX-H3K9me3 | 10 | H3K27ac-cFos | 15 | H3K9me2-SUV39H1 | 16 | H3S10ph-PIM1 | 16 |
| H3K9me3-MSK2 | 10 | H3K27ac-cJun | 15 | Aurora_B-MSK2 | 16 | H3K79me3-KMT2B | 16 |
| H3K9me3-Myc | 10 | H3K27ac-PIM1 | 15 | H3K9me2-SETD2 | 16 | CDK8-EHMT2 | 16 |
| H3K9me2-H3K9me2 | 10 | H3K27me3-KMT2A | 16 | CBP-YY1 | 16 | EHMT1-EP300 | 16 |
| EHMT2-H3K9me3 | 10 | H3K9me3-NRF1 | 16 | H3K79me3-SETD2 | 16 | Aurora_B-cFos | 16 |
| H3K9me3-H3S10ph | 10 | H3K9me3-PIM1 | 16 | EHMT2-SETD2 | 16 | Aurora_B-KMT2A | 16 |
| H3K27me3-H3K9me3 | 11 | H3K9me3-MSK1 | 16 | Aurora_B-H3K36me3 | 16 | EHMT2-SUV39H1 | 16 |
| H3K27me3-H3K4me1 | 11 | H3K79me3-H3K9me3 | 16 | Aurora_B-H3K14ac | 16 | KMT2A-cJun | 16 |
| H3K27me3-H3K27me3 | 11 | EZH2-H3K9me3 | 16 | H3K14ac-SETD2 | 16 | MSK2-SUV39H1 | 16 |
| Aurora_B-H3K27me3 | 11 | H3K36me3-H3K9me3 | 16 | Myc-SETD2 | 16 | MSK2-MSK2 | 16 |
| H3K27me3-KMT2B | 11 | H3K14ac-H3K9me3 | 16 | KMT2B-YY1 | 16 | CTCF-H3S10ph | 16 |
| H3K27me3-SUV39H1 | 11 | H3K9me3-KMT2A | 16 | KMT2B-USF1 | 16 | CDK8-USF2 | 16 |
| CBP-H3K27me3 | 11 | H3K9ac-MSK2 | 16 | SETD2-cFos | 16 | EZH2-SETD2 | 16 |
| CDK8-H3K27me3 | 12 | H3K4me1-USF1 | 16 | SETD2-USF2 | 16 | EHMT2-EHMT2 | 16 |
| CTCF-H3K27me3 | 12 | H3K14ac-H3K27ac | 16 | H3K9me2-KMT2A | 16 | CTCF-PIM1 | 16 |

| | | | | | | | |
|---|---|---|---|---|---|---|---|
| H3K27me3-SETD2 | 12 | H3K27ac-USF2 | 16 | CTCF-KMT2A | 16 | H3K9ac-USF1 | 16 |
| H3K27me3-H3K9me2 | 12 | H3K27ac-Myc | 16 | CTCF-EZH2 | 16 | CTCF-H3K36me3 | 16 |
| H3K27me3-H3K36me3 | 12 | cJun-cJun | 16 | γH2AX-γH2AX | 16 | EP300-cFos | 16 |
| H3K27me3-cJun | 12 | Aurora_B-KMT2B | 16 | Aurora_B-USF2 | 16 | KMT2A-SETD2 | 16 |
| EZH2-H3K27me3 | 12 | Aurora_B-SUV39H1 | 16 | CBP-H3K9me2 | 16 | EP300-USF2 | 16 |
| H3K27me3-MSK2 | 12 | H3K27me3-cFos | 16 | KMT2B-USF2 | 16 | PIM1-PIM1 | 16 |
| H3K27me3-H3S10ph | 12 | H3K27me3-USF2 | 16 | H3S10ph-cJun | 16 | PIM1-SETD2 | 16 |
| H3K27me3-PIM1 | 12 | H3K27me3-NRF1 | 16 | EHMT1-SUV39H1 | 16 | CBP-H3S10ph | 16 |
| H3K27me3-H3K79me3 | 12 | EHMT1-H3K27me3 | 16 | NRF1-SUV39H1 | 16 | H3S10ph-SUV39H1 | 16 |
| H3K14ac-H3K27me3 | 12 | Aurora_B-CDK8 | 16 | CBP-NRF1 | 16 | CBP-MSK2 | 16 |
| H3K27me3-MSK1 | 12 | H3K27me3-Myc | 16 | KMT2B-NRF1 | 16 | MSK2-SETD2 | 16 |
| EHMT2-H3K27me3 | 12 | H3K9ac-H3S10ph | 16 | γH2AX-H3K9me2 | 16 | MSK1-SETD2 | 16 |
| Aurora_B-Aurora_B | 12 | EP300-cJun | 16 | CTCF-γH2AX | 16 | CDK8-H3K14ac | 16 |
| γH2AX-H3K27me3 | 12 | H3K14ac-H3K9ac | 16 | CBP-EHMT1 | 16 | NRF1-NRF1 | 16 |
| H3K4me1-SUV39H1 | 13 | CDK8-SETD2 | 16 | EHMT1-KMT2B | 16 | CDK8-γH2AX | 16 |
| CBP-H3K4me1 | 13 | CBP-MSK1 | 16 | MSK2-cJun | 16 | CDK8-MSK2 | 16 |
| H3K4me1-KMT2B | 13 | Aurora_B-EP300 | 16 | EHMT2-EP300 | 16 | EP300-H3K14ac | 16 |

| | | | | | | | |
|---|---|---|---|---|---|---|---|
| CTCF-H3K4me1 | 13 | CDK8-H3K9me2 | 16 | CBP-Myc | 16 | EP300-EZH2 | 16 |
| H3K4me1-H3K9me2 | 13 | CDK8-CTCF | 16 | H3K36me3-cJun | 16 | CDK8-EZH2 | 16 |
| H3K4me1-SETD2 | 13 | CBP-cJun | 16 | H3K9me2-PIM1 | 16 | CDK8-cJun | 16 |
| H3K27me3-H3K9ac | 13 | SUV39H1-cJun | 16 | γH2AX-cJun | 16 | EP300-H3K36me3 | 16 |
| EP300-H3K27me3 | 13 | EZH2-EZH2 | 16 | H3K9me2-H3S10ph | 16 | Max-RNAPII | 16 |
| Aurora_B-H3K4me1 | 13 | EZH2-SUV39H1 | 16 | H3K9me2-Myc | 16 | EHMT1-H3K9ac | 16 |
| EP300-H3K9me3 | 14 | H3K9ac-NRF1 | 16 | CTCF-Myc | 16 | EP300-MSK1 | 16 |
| CDK8-H3K9me3 | 14 | KMT2A-SUV39H1 | 16 | H3K79me3-H3K79me3 | 16 | EP300-H3S10ph | 16 |
| EP300-H3K27ac | 14 | KMT2B-SETD2 | 16 | H3K9me2-MSK2 | 16 | EHMT1-H3K27ac | 16 |
| H3K27ac-SUV39H1 | 14 | CBP-SETD2 | 16 | MSK1-MSK2 | 16 | EHMT2-H3K9ac | 16 |
| H3K9ac-H3K9ac | 14 | SETD2-SUV39H1 | 16 | Myc-cJun | 16 | EP300-KMT2A | 16 |
| H3K9ac-SUV39H1 | 14 | CBP-KMT2B | 16 | γH2AX-H3S10ph | 16 | H3K9ac-USF2 | 16 |
| EP300-H3K9ac | 14 | KMT2B-SUV39H1 | 16 | Myc-Myc | 16 | EP300-PIM1 | 16 |
| H3K27ac-KMT2B | 14 | Aurora_B-CBP | 16 | γH2AX-H3K14ac | 16 | NRF1-RNAPII | 16 |
| CBP-H3K27ac | 14 | EHMT1-H3K9me3 | 16 | CTCF-EHMT2 | 16 | EZH2-H3K9ac | 16 |
| CDK8-H3K27ac | 14 | H3K9me3-USF2 | 16 | CDK8-YY1 | 16 | H3K9ac-MSK1 | 16 |

| | | | | | | | |
|---|---|---|---|---|---|---|---|
| EP300-EP300 | 14 | KMT2B-cJun | 16 | H3K36me3-H3K79me3 | 16 | KMT2B-PIM1 | 16 |
| H3K27ac-H3K9me2 | 14 | CBP-CTCF | 16 | CTCF-H3K79me3 | 16 | H3K9ac-Myc | 16 |
| CTCF-H3K27ac | 14 | KMT2A-KMT2B | 16 | EZH2-PIM1 | 16 | EHMT2-H3K27ac | 16 |
| H3K27ac-SETD2 | 14 | MSK1-MSK1 | 16 | PIM1-cJun | 16 | EZH2-H3K27ac | 16 |
| H3K4me1-cJun | 15 | CDK8-KMT2A | 16 | MSK1-cJun | 16 | H3K27ac-H3K36me3 | 16 |
| CBP-CBP | 15 | CDK8-Myc | 16 | Aurora_B-Myc | 16 | H3K79me3-H3K9ac | 16 |
| PIM1-RNAPII | 15 | RNAPII-USF1 | 16 | Aurora_B-EHMT2 | 16 | H3K9ac-cFos | 16 |
| H3S10ph-RNAPII | 15 | CDK8-PIM1 | 16 | H3K14ac-H3K14ac | 16 | | |
| CTCF-RNAPII | 15 | CDK8-H3K79me3 | 16 | CTCF-MSK2 | 16 | | |
| Aurora_B-RNAPII | 15 | SETD2-cJun | 16 | CTCF-H3K9me2 | 16 | | |

**Table S3. Antibody**

| # | Target | Vendor | Catalog |
|---|---|---|---|
| C | IgG_control | abcam | ab37415 |
| 1 | H3K36me3 | CST | 4909BF |
| 2 | H3K4me1 | CST | 5326BF |
| 3 | H3K27ac | CST | 8173BF |
| 4 | H3S10ph | abcam | ab239405 |
| 5 | gH2AX | abcam | ab215967 |
| 6 | H3K79me3 | CST | 74073BF |
| 7 | H3K9me2 | CST | 4658BF |
| 8 | H3K9me3 | abcam | ab232324 |

| | | | |
|---|---|---|---|
| 9 | H3K14ac | CST | 7627BF |
| 10 | H3K27me3 | CST | 9733BF |
| 11 | H3K4me3 | CST | 9751BF |
| 12 | SETD2 | CST | 80290BF |
| 13 | KMT2B | CST | 63735BF |
| 14 | CBP | CST | 7389BF |
| 15 | EP300 | abcam | ab275388 |
| 16 | MSK1 | CST | 3489BF |
| 17 | MSK2 | CST | 3679BF |
| 18 | PIM1 | CST | 54523BF |
| 19 | CDK8 | CST | 17395BF |
| 20 | Aurora_B | CST | 28711BF |
| 21 | EHMT2 | CST | 68851BF |
| 22 | SUV39H1 | CST | 8729BF |
| 23 | EHMT1 | CST | 35005BF |
| 24 | EZH2 | CST | 5246BF |
| 25 | KMT2A | CST | 14689BF |
| 26 | CTCF | CST | 3418BF |
| 27 | RNAPII | CST | 13499BF |
| 28 | cJun | abcam | ab218576 |
| 29 | cFos | CDI | 20221011 |
| 30 | Max | CDI | 20220329 |
| 31 | Myc | abcam | ab168727 |
| 32 | USF1 | CDI | 20221025 |
| 33 | USF2 | CDI | 20221010 |
| 34 | NRF1 | CDI | 20221004 |
| 35 | YY1 | CDI | 20221011 |
| 36 | H3K9ac | abcam | ab203951 |

**Table S4. Reagents**

| Product name | Vendor | Catalog |
|---|---|---|
| **Antibody Barcoding** | | |
| EZ-Link NHS-PEG12-Biotin, No-Weigh Format | ThermoFisher | A35389 |
| Zeba™ Spin Desalting Columns, 40K MWCO, 0.5 mL | ThermoFisher | 87767 |
| Zeba™ 96-well Spin Desalting Plates, 40K MWCO | ThermoFisher | 87775 |
| Streptavidin Protein | ThermoFisher | 21122 |
| D-biotin | ThermoFisher | B20656 |
| Amicon Ultra 0.5 Centrifugal Filter 30 kDa MWCO | Millipore Sigma | UFC503096 |
| AMICON ULTRA-4 CENTRIFUGAL FILTER UNIT WITH ULTRACEL-30 MEMBRANE | MilliporeSigma | UFC803096 |
| UltraPure DNase/RNase-Free Distilled Water | ThermoFisher | 10977023 |
| **Hi-Plex CUT&Tag** | | |
| BioMag®Plus Concanavalin A | Polysciences | 86057-3 |
| HEPES (1 M) | ThermoFisher | 15630080 |
| NaCl (5 M), RNase-free | ThermoFisher | AM9760G |
| KCl (2 M), RNase-free | ThermoFisher | AM9640G |
| SPERMIDINE 0.1 M SOLUTION | MilliporeSigma | 05292-1ML-F |
| Digitonin (5%) | ThermoFisher | BN2006 |
| CALCIUM CHLORIDE SOLUTION | MilliporeSigma | 21115-100ML |
| MANGANESE(II) CHLORIDE SOLUTION | MilliporeSigma | M1787-100ML |
| cOmplete(TM), EDTA-free Protease Inhibitor Cocktail | MilliporeSigma | 11873580001 |
| Tagmentase | Diagenode | C01070010-20 |
| MgCl^2^ (1 M) | ThermoFisher | AM9530G |
| Corning(R) 100 mL 0.5M EDTA, pH 8.0 | Corning Cellgro | 46-034-CI |
| Corning(R) 100 mL SDS (Sodium Dodecyl Sulfate) | Corning Cellgro | 46-040-CI |
| Proteinase K, recombinant, PCR grade | ThermoFisher | EO0491 |
| Phenol:Chloroform + Tris Buffer | ThermoFisher | 17908 |
| 5Prime Phase Lock Gel Heavy 200 x 2 mL | ThermoFisher | NC1093153 |
| GLYCOGEN MB GRADE | ThermoFisher | R0561 |
| RNase A, DNase and protease-free (10 mg/mL) | ThermoFisher | EN0531 |
| TRIS HCl, 1M pH 8.0 500ml | QualityBiological | 351-007-101 |

| | | |
|---|---|---|
| NEBNext Ultra II Q5 Master Mix - 50 reactions | NEB | M0544S |
| AMPure XP Reagent, 60 mL | Beckman | A63881 |
| **scHi-Plex CUT&Tag** | | |
| TERGITOL TYPE NP-40 70% IN H2O | MilliporeSigma | NP40S-500ML |
| Triton™ X-100 | MilliporeSigma | T8787-100ml |
| DAPI | ThermoFisher | D1306 |
| Falcon® 5 mL Round Bottom Polystyrene Test Tube, with Cell Strainer Snap Cap | Falcon | 352235 |
| Single-well Deep Well Plates | Miltenyi Biotec | 130-114-966 |
| **Cell culture** | | |
| RPMI 1640 Medium | Gibco | 11875119 |
| FetalPlex animal serum complex | Gemini Bio | 100-602-500 |
| Penicillin-Streptomycin (10,000 U/mL) | ThermoFisher | 15140122 |
| SODIUM BUTYRATE 10ML | MilliporeSigma | 19-137 |
| **RNA-seq preparation** | | |
| miRNeasy Mini Kit (50) | Qiagen | 217004 |
| UltraPure™ DEPC-Treated Water | ThermoFisher | 750024 |
| Dynabeads™ mRNA Purification Kit | Invitrogen | 61006 |
| NEBNext® Ultra™ II RNA Library Prep Kit for Illumina | NEB | E7770S |
| NEBNext® Multiplex Oligos for Illumina® (Index Primers Set 1) | NEB | E7335S |

**Table S5. Barcode assignment and adaptor sequence**

| # | Target | Barcode | Name | Sequence |
|---|---|---|---|---|
| **P5 adaptor** | | | | |
| C | IgG_ control | AGTGCCCTAGA | 11nt_Bio-P5-C | /5Biosg/TCGTCGGCAGCGTCTCCACGCAGTGCCCTAGAGCGATCGAGGACGGCAGATGTGTATAAGAGACAG |
| 1 | H3K36me3 | GTCTATGCGTT | 11nt_Bio-P5-1 | /5Biosg/TCGTCGGCAGCGTCTCCACGCGTCTATGCGTTGCGATCGAGGACGGCAGATGTGTATAAGAGACAG |
| 2 | H3K4me1 | ATTTCCGGTCG | 11nt_Bio-P5-2 | /5Biosg/TCGTCGGCAGCGTCTCCACGCATTTCCGGTCGGCGATCGAGGACGGCAGATGTGTATAAGAGACAG |

| | | | | |
|---|---|---|---|---|
| 3 | H3K27ac | CAAACGTGAGG | 11nt_Bio-P5-3 | /5Biosg/TCGTCGGCAGCGTCTCCACGCCAAACGTGAGGGCGATCGAGGACGGCAGATGTGTATAAGAGACAG |
| 4 | H3S10ph | CCTCCAACAAT | 11nt_Bio-P5-4 | /5Biosg/TCGTCGGCAGCGTCTCCACGCCCTCCAACAATGCGATCGAGGACGGCAGATGTGTATAAGAGACAG |
| 5 | gH2AX | GGCTTATGCAC | 11nt_Bio-P5-5 | /5Biosg/TCGTCGGCAGCGTCTCCACGCGGCTTATGCACGCGATCGAGGACGGCAGATGTGTATAAGAGACAG |
| 6 | H3K79me3 | GGTAGTCCTGT | 11nt_Bio-P5-6 | /5Biosg/TCGTCGGCAGCGTCTCCACGCGGTAGTCCTGTGCGATCGAGGACGGCAGATGTGTATAAGAGACAG |
| 7 | H3K9me2 | CGGAGCCTAAT | 11nt_Bio-P5-7 | /5Biosg/TCGTCGGCAGCGTCTCCACGCCGGAGCCTAATGCGATCGAGGACGGCAGATGTGTATAAGAGACAG |
| 8 | H3K9me3 | TAGGTGCAAAG | 11nt_Bio-P5-8 | /5Biosg/TCGTCGGCAGCGTCTCCACGCTAGGTGCAAAGGCGATCGAGGACGGCAGATGTGTATAAGAGACAG |
| 9 | H3K14ac | TTGGAGTTGCA | 11nt_Bio-P5-9 | /5Biosg/TCGTCGGCAGCGTCTCCACGCTTGGAGTTGCAGCGATCGAGGACGGCAGATGTGTATAAGAGACAG |
| 10 | H3K27me3 | TTTGACGGTTA | 11nt_Bio-P5-10 | /5Biosg/TCGTCGGCAGCGTCTCCACGCTTTGACGGTTAGCGATCGAGGACGGCAGATGTGTATAAGAGACAG |
| 11 | H3K4me3 | CGCGGGTATAT | 11nt_Bio-P5-11 | /5Biosg/TCGTCGGCAGCGTCTCCACGCCGCGGGTATATGCGATCGAGGACGGCAGATGTGTATAAGAGACAG |
| 12 | SETD2 | GCCCATTAAAT | 11nt_Bio-P5-12 | /5Biosg/TCGTCGGCAGCGTCTCCACGCGCCCATTAAATGCGATCGAGGACGGCAGATGTGTATAAGAGACAG |
| 13 | KMT2B | TCCATCTTAAG | 11nt_Bio-P5-13_2 | /5Biosg/TCGTCGGCAGCGTCTCCACGCTCCATCTTAAGGCGATCGAGGACGGCAGATGTGTATAAGAGACAG |
| 14 | CBP | TAAGTAAGCCT | 11nt_Bio-P5-14 | /5Biosg/TCGTCGGCAGCGTCTCCACGCTAAGTAAGCCTGCGATCGAGGACGGCAGATGTGTATAAGAGACAG |
| 15 | EP300 | ATACTCCCACT | 11nt_Bio-P5-15 | /5Biosg/TCGTCGGCAGCGTCTCCACGCATACTCCCACTGCGATCGAGGACGGCAGATGTGTATAAGAGACAG |
| 16 | MSK1 | GTACCGGGTTA | 11nt_Bio-P5-16 | /5Biosg/TCGTCGGCAGCGTCTCCACGCGTACCGGGTTAGCGATCGAGGACGGCAGATGTGTATAAGAGACAG |
| 17 | MSK2 | GGATCATTTAG | 11nt_Bio-P5-17 | /5Biosg/TCGTCGGCAGCGTCTCCACGCGGATCATTTAGGCGATCGAGGACGGCAGATGTGTATAAGAGACAG |
| 18 | PIM1 | TTAAACCCGTC | 11nt_Bio-P5-18 | /5Biosg/TCGTCGGCAGCGTCTCCACGCTTAAACCCGTCGCGATCGAGGACGGCAGATGTGTATAAGAGACAG |

| | | | | |
|---|---|---|---|---|
| 19 | CDK8 | CCGGAA ATCAC | 11nt_Bio-P5-19 | /5Biosg/TCGTCGGCAGCGTCTCCACGCCCGGAAATCA CGCGATCGAGGACGGCAGATGTGTATAAGAGACAG |
| 20 | Aurora_B | TCTCAT CGGCT | 11nt_Bio-P5-20 | /5Biosg/TCGTCGGCAGCGTCTCCACGCTCTCATCGGC TGCGATCGAGGACGGCAGATGTGTATAAGAGACAG |
| 21 | EHMT2 | AGAGCG TCATT | 11nt_Bio-P5-21 | /5Biosg/TCGTCGGCAGCGTCTCCACGCAGAGCGTCAT TGCGATCGAGGACGGCAGATGTGTATAAGAGACAG |
| 22 | SUV39H1 | TCCTAG CCTAC | 11nt_Bio-P5-22 | /5Biosg/TCGTCGGCAGCGTCTCCACGCTCCTAGCCTA CGCGATCGAGGACGGCAGATGTGTATAAGAGACAG |
| 23 | EHMT1 | CGAACC AACCA | 11nt_Bio-P5-23 | /5Biosg/TCGTCGGCAGCGTCTCCACGCCGAACCAACC AGCGATCGAGGACGGCAGATGTGTATAAGAGACAG |
| 24 | EZH2 | AGATAG CAGTC | 11nt_Bio-P5-24 | /5Biosg/TCGTCGGCAGCGTCTCCACGCAGATAGCAGT CGCGATCGAGGACGGCAGATGTGTATAAGAGACAG |
| 25 | KMT2A | AGTCCG AACTC | 11nt_Bio-P5-25 | /5Biosg/TCGTCGGCAGCGTCTCCACGCAGTCCGAACT CGCGATCGAGGACGGCAGATGTGTATAAGAGACAG |
| 26 | CTCF | AGTATT TCGCG | 11nt_Bio-P5-26 | /5Biosg/TCGTCGGCAGCGTCTCCACGCAGTATTTCGC GGCGATCGAGGACGGCAGATGTGTATAAGAGACAG |
| 27 | RNAPII | CTACAA AGCCG | 11nt_Bio-P5-27 | /5Biosg/TCGTCGGCAGCGTCTCCACGCCTACAAAGCC GGCGATCGAGGACGGCAGATGTGTATAAGAGACAG |
| 28 | cJun | ACTACG CATCT | 11nt_Bio-P5-28 | /5Biosg/TCGTCGGCAGCGTCTCCACGCACTACGCATCT GCGATCGAGGACGGCAGATGTGTATAAGAGACAG |
| 29 | cFos | ATTGCC AACCT | 11nt_Bio-P5-29 | /5Biosg/TCGTCGGCAGCGTCTCCACGCATTGCCAACCT GCGATCGAGGACGGCAGATGTGTATAAGAGACAG |
| 30 | Max | ACCCGT AAAGG | 11nt_Bio-P5-30 | /5Biosg/TCGTCGGCAGCGTCTCCACGCACCCGTAAAG GGCGATCGAGGACGGCAGATGTGTATAAGAGACAG |
| 31 | Myc | CCGTGC ACTTT | 11nt_Bio-P5-31 | /5Biosg/TCGTCGGCAGCGTCTCCACGCCCGTGCACTT TGCGATCGAGGACGGCAGATGTGTATAAGAGACAG |
| 32 | USF1 | AGCCCA ATCGA | 11nt_Bio-P5-32 | /5Biosg/TCGTCGGCAGCGTCTCCACGCAGCCCAATCG AGCGATCGAGGACGGCAGATGTGTATAAGAGACAG |
| 33 | USF2 | CCTATT AGGAG | 11nt_Bio-P5-33 | /5Biosg/TCGTCGGCAGCGTCTCCACGCCCTATTAGGA GGCGATCGAGGACGGCAGATGTGTATAAGAGACAG |
| 34 | NRF1 | ATAGTC GAATG | 11nt_Bio-P5-34 | /5Biosg/TCGTCGGCAGCGTCTCCACGCATAGTCGAAT GGCGATCGAGGACGGCAGATGTGTATAAGAGACAG |

| | | | | |
|---|---|---|---|---|
| 35 | YY1 | TACTGT AGGTC | 11nt_Bio-P5-35 | /5Biosg/TCGTCGGCAGCGTCTCCACGCTACTGTAGGTCGCGATCGAGGACGGCAGATGTGTATAAGAGACAG |
| 36 | H3K9ac | ACGCTA CTCTT | 11nt_Bio-P5-36 | /5Biosg/TCGTCGGCAGCGTCTCCACGCACGCTACTCTTGCGATCGAGGACGGCAGATGTGTATAAGAGACAG |
| **P7 adaptor** | | | | |
| C | IgG_ control | AGTGCC CTAGA | 11nt_Bio-P7-C | /5Biosg/GTCTCGTGGGCTCGGCTGTCCCTGTCCAGTGCCCTAGACACCGTCTCCGCCTCAGATGTGTATAAGAGACAG |
| 1 | H3K36me3 | GTCTAT GCGTT | 11nt_Bio-P7-1 | /5Biosg/GTCTCGTGGGCTCGGCTGTCCCTGTCCGTCTATGCGTTCACCGTCTCCGCCTCAGATGTGTATAAGAGACAG |
| 2 | H3K4me1 | ATTTCC GGTCG | 11nt_Bio-P7-2 | /5Biosg/GTCTCGTGGGCTCGGCTGTCCCTGTCCATTTCCGGTCGCACCGTCTCCGCCTCAGATGTGTATAAGAGACAG |
| 3 | H3K27ac | CAAACG TGAGG | 11nt_Bio-P7-3 | /5Biosg/GTCTCGTGGGCTCGGCTGTCCCTGTCCCAAACGTGAGGCACCGTCTCCGCCTCAGATGTGTATAAGAGACAG |
| 4 | H3S10ph | CCTCCA ACAAT | 11nt_Bio-P7-4 | /5Biosg/GTCTCGTGGGCTCGGCTGTCCCTGTCCCCTCCAACAATCACCGTCTCCGCCTCAGATGTGTATAAGAGACAG |
| 5 | gH2AX | GGCTTA TGCAC | 11nt_Bio-P7-5 | /5Biosg/GTCTCGTGGGCTCGGCTGTCCCTGTCCGGCTTATGCACCACCGTCTCCGCCTCAGATGTGTATAAGAGACAG |
| 6 | H3K79me3 | GGTAGT CCTGT | 11nt_Bio-P7-6 | /5Biosg/GTCTCGTGGGCTCGGCTGTCCCTGTCCGGTAGTCCTGTCACCGTCTCCGCCTCAGATGTGTATAAGAGACAG |
| 7 | H3K9me2 | CGGAG CCTAAT | 11nt_Bio-P7-7 | /5Biosg/GTCTCGTGGGCTCGGCTGTCCCTGTCCCGGAGCCTAATCACCGTCTCCGCCTCAGATGTGTATAAGAGACAG |
| 8 | H3K9me3 | TAGGTG CAAAG | 11nt_Bio-P7-8 | /5Biosg/GTCTCGTGGGCTCGGCTGTCCCTGTCCTAGGTGCAAAGCACCGTCTCCGCCTCAGATGTGTATAAGAGACAG |

| | | | | |
|---|---|---|---|---|
| 9 | H3K14ac | TTGGAG TTGCA | 11nt_Bio-P7-9 | /5Biosg/GTCTCGTGGGCTCGGCTGTCCCTGTCCTTGGAGTTGCACACCGTCTCCGCCTCAGATGTGTATAAGAGACAG |
| 10 | H3K27me3 | TTTGAC GGTTA | 11nt_Bio-P7-10 | /5Biosg/GTCTCGTGGGCTCGGCTGTCCCTGTCCTTTGACGGTTACACCGTCTCCGCCTCAGATGTGTATAAGAGACAG |
| 11 | H3K4me3 | CGCGG GTATAT | 11nt_Bio-P7-11 | /5Biosg/GTCTCGTGGGCTCGGCTGTCCCTGTCCCGCGGGTATATCACCGTCTCCGCCTCAGATGTGTATAAGAGACAG |
| 12 | SETD2 | GCCCAT TAAAT | 11nt_Bio-P7-12 | /5Biosg/GTCTCGTGGGCTCGGCTGTCCCTGTCCGCCCATTAAATCACCGTCTCCGCCTCAGATGTGTATAAGAGACAG |
| 13 | KMT2B | TCCATC TTAAG | 11nt_Bio-P7-13_2 | /5Biosg/GTCTCGTGGGCTCGGCTGTCCCTGTCCTCCATCTTAAGCACCGTCTCCGCCTCAGATGTGTATAAGAGACAG |
| 14 | CBP | TAAGTA AGCCT | 11nt_Bio-P7-14 | /5Biosg/GTCTCGTGGGCTCGGCTGTCCCTGTCCTAAGTAAGCCTCACCGTCTCCGCCTCAGATGTGTATAAGAGACAG |
| 15 | EP300 | ATACTC CCACT | 11nt_Bio-P7-15 | /5Biosg/GTCTCGTGGGCTCGGCTGTCCCTGTCCATACTCCCACTCACCGTCTCCGCCTCAGATGTGTATAAGAGACAG |
| 16 | MSK1 | GTACCG GGTTA | 11nt_Bio-P7-16 | /5Biosg/GTCTCGTGGGCTCGGCTGTCCCTGTCCGTACCGGGTTACACCGTCTCCGCCTCAGATGTGTATAAGAGACAG |
| 17 | MSK2 | GGATCA TTTAG | 11nt_Bio-P7-17 | /5Biosg/GTCTCGTGGGCTCGGCTGTCCCTGTCCGGATCATTTAGCACCGTCTCCGCCTCAGATGTGTATAAGAGACAG |
| 18 | PIM1 | TTAAAC CCGTC | 11nt_Bio-P7-18 | /5Biosg/GTCTCGTGGGCTCGGCTGTCCCTGTCCTTAAACCCGTCCACCGTCTCCGCCTCAGATGTGTATAAGAGACAG |
| 19 | CDK8 | CCGGAA ATCAC | 11nt_Bio-P7-19 | /5Biosg/GTCTCGTGGGCTCGGCTGTCCCTGTCCCCGGAAATCACCACCGTCTCCGCCTCAGATGTGTATAAGAGACAG |

| | | | | |
|---|---|---|---|---|
| 20 | Aurora_B | TCTCAT CGGCT | 11nt_Bio-P7-20 | /5Biosg/GTCTCGTGGGCTCGGCTGTCCCTGTCCTCTCATCGGCTCACCGTCTCCGCCTCAGATGTGTATAAGAGACAG |
| 21 | EHMT2 | AGAGCG TCATT | 11nt_Bio-P7-21 | /5Biosg/GTCTCGTGGGCTCGGCTGTCCCTGTCCAGAGCGTCATTCACCGTCTCCGCCTCAGATGTGTATAAGAGACAG |
| 22 | SUV39H1 | TCCTAG CCTAC | 11nt_Bio-P7-22 | /5Biosg/GTCTCGTGGGCTCGGCTGTCCCTGTCCTCCTAGCCTACCACCGTCTCCGCCTCAGATGTGTATAAGAGACAG |
| 23 | EHMT1 | CGAACC AACCA | 11nt_Bio-P7-23 | /5Biosg/GTCTCGTGGGCTCGGCTGTCCCTGTCCCGAACCAACCACACCGTCTCCGCCTCAGATGTGTATAAGAGACAG |
| 24 | EZH2 | AGATAG CAGTC | 11nt_Bio-P7-24 | /5Biosg/GTCTCGTGGGCTCGGCTGTCCCTGTCCAGATAGCAGTCCACCGTCTCCGCCTCAGATGTGTATAAGAGACAG |
| 25 | KMT2A | AGTCCG AACTC | 11nt_Bio-P7-25 | /5Biosg/GTCTCGTGGGCTCGGCTGTCCCTGTCCAGTCCGAACTCCACCGTCTCCGCCTCAGATGTGTATAAGAGACAG |
| 26 | CTCF | AGTATT TCGCG | 11nt_Bio-P7-26 | /5Biosg/GTCTCGTGGGCTCGGCTGTCCCTGTCCAGTATTTCGCGCACCGTCTCCGCCTCAGATGTGTATAAGAGACAG |
| 27 | RNAPII | CTACAA AGCCG | 11nt_Bio-P7-27 | /5Biosg/GTCTCGTGGGCTCGGCTGTCCCTGTCCCTACAAAGCCGCACCGTCTCCGCCTCAGATGTGTATAAGAGACAG |
| 28 | cJun | ACTACG CATCT | 11nt_Bio-P7-28 | /5Biosg/GTCTCGTGGGCTCGGCTGTCCCTGTCCACTACGCATCTCACCGTCTCCGCCTCAGATGTGTATAAGAGACAG |
| 29 | cFos | ATTGCC AACCT | 11nt_Bio-P7-29 | /5Biosg/GTCTCGTGGGCTCGGCTGTCCCTGTCCATTGCCAACCTCACCGTCTCCGCCTCAGATGTGTATAAGAGACAG |
| 30 | Max | ACCCGT AAAGG | 11nt_Bio-P7-30 | /5Biosg/GTCTCGTGGGCTCGGCTGTCCCTGTCCACCCGTAAAGGCACCGTCTCCGCCTCAGATGTGTATAAGAGACAG |

| | | | | |
|---|---|---|---|---|
| 31 | Myc | CCGTGCACTTT | 11nt_Bio-P7-31 | /5Biosg/GTCTCGTGGGCTCGGCTGTCCCTGTCCCCGTGCACTTTCACCGTCTCCGCCTCAGATGTGTATAAGAGACAG |
| 32 | USF1 | AGCCCAATCGA | 11nt_Bio-P7-32 | /5Biosg/GTCTCGTGGGCTCGGCTGTCCCTGTCCAGCCCAATCGACACCGTCTCCGCCTCAGATGTGTATAAGAGACAG |
| 33 | USF2 | CCTATTAGGAG | 11nt_Bio-P7-33 | /5Biosg/GTCTCGTGGGCTCGGCTGTCCCTGTCCCCTATTAGGAGCACCGTCTCCGCCTCAGATGTGTATAAGAGACAG |
| 34 | NRF1 | ATAGTCGAATG | 11nt_Bio-P7-34 | /5Biosg/GTCTCGTGGGCTCGGCTGTCCCTGTCCATAGTCGAATGCACCGTCTCCGCCTCAGATGTGTATAAGAGACAG |
| 35 | YY1 | TACTGTAGGTC | 11nt_Bio-P7-35 | /5Biosg/GTCTCGTGGGCTCGGCTGTCCCTGTCCTACTGTAGGTCCACCGTCTCCGCCTCAGATGTGTATAAGAGACAG |
| 36 | H3K9ac | ACGCTACTCTT | 11nt_Bio-P7-36 | /5Biosg/GTCTCGTGGGCTCGGCTGTCCCTGTCCACGCTACTCTTCACCGTCTCCGCCTCAGATGTGTATAAGAGACAG |

**Table S6. Oligo sequence**

| Name | Function | Sequence |
|---|---|---|
| Tn5MErev | Reverse Primer anneal with connector for loading to Tn5 | [phos]CTGTCTCTTATACACATCT |
| MCT Read1 | Custom read1 primer for P5-BC sequencing | TCGTCGGCAGCGTCTCCACGC |
| MCT Read2 | Custom read2 primer for P7-BC sequencing | GTCTCGTGGGCTCGGCTGTCCCTGTCC |
| MCT Index1 | Custom Index1 primer for i7 sequencing | GGACAGGGACAGCCGAGCCCACGAGAC |
| MCT Index2 | Custom Index2 primer for i5 sequencing | GCGTGGAGACGCTGCCGACGA |

### REFERENCES

1. Love, M.I., W. Huber, and S. Anders, *Moderated estimation of fold change and dispersion for RNA-seq data with DESeq2.* Genome Biol, 2014. **15**(12): p. 550.
2. H, W., *ggplot2: Elegant Graphics for Data Analysis.* Springer-Verlag New York. , 2016. **ISBN 978-3-319-24277-4**.
